# Wash-Free Multi-Target Super-Resolution Microscopy with Photocaged DNA Labels

**DOI:** 10.64898/2026.01.13.699198

**Authors:** Nina Kaltenschnee, Marina S. Dietz, Laurell F. Kessler, Yunqing Li, Janik Kaufmann, Alexander Heckel, Mike Heilemann

**Affiliations:** Institute of Organic Chemistry and Chemical Biology, Goethe University Frankfurt, Frankfurt am Main, Germany; Institute of Physical and Theoretical Chemistry, Goethe University Frankfurt, Frankfurt am Main, Germany

**Author notes:** These authors contributed equally (Nina Kaltenschnee, Marina S. Dietz).

**Keywords:** DNA-PAINT, light control, multiplexed imaging, photocages, single-molecule localization microscopy

## Abstract

Super-resolution microscopy with DNA-fluorophore labels is primed for multi-target imaging of cell biological samples. However, direct interaction with the sample is required to exchange or add DNA-fluorophore labels in each imaging round, which can impair the accuracy of the imaging data at the nanometer scale. To bypass this requirement, we introduce a wash-free method that employs DNA oligonucleotides equipped with photocaging groups. Irradiation with light removes these photo-modulatable groups and changes the hybridization properties of DNA labels, enabling light-modulated targeting. We demonstrate this concept by imaging various cellular targets with confocal microscopy, single-molecule localization microscopy, and stimulated emission depletion (STED) microscopy.

## Introduction

Super-resolution microscopy has revolutionized our understanding of cell biology.^[1]^ To capture the full context of subcellular architecture, imaging multiple targets within a sample is essential. Given that only a limited number of fluorophores can be distinguished based on their optical properties, this limitation has driven the development of sequential imaging strategies to increase the number of accessible targets in an experiment. This can, for example, be realized with chemical strategies that remove and replace fluorophore labels, as demonstrated in single-molecule localization microscopy.^[2,3]^ Alternatively, fluorophore labels with transient, weak binding affinities to their target allow multi-target imaging through iterative exchange steps coupled with washing.^[4]^ Examples are short, transiently hybridizing and fluorophore-labeled DNA oligonucleotides, as in DNA-point accumulation for nanoscale topography (DNA-PAINT),^[5]^ weak-affinity fluorophore labels for orthogonal self-labeling proteins,^[6,7]^ and non-covalent small-molecule fluorophore labels.^[8]^ Notably, these labeling strategies are compatible with various modalities of super-resolution microscopy.^[6,9–11]^

Fluorophore-labeled DNA in combination with sequence design has demonstrated many-target fluorescence imaging of complex molecular samples.^[12,13]^ One remaining challenge of these strategies is that either washing steps are needed, or molecular components need to be added to and mixed with the imaging buffer. Such direct interaction with the sample introduces an additional step in the experimental workflow and may perturb the nanoscale architecture of cellular samples. This, in turn, affects the information content that would otherwise be accessible with the excellent spatial resolution that these methods can achieve.

To avoid physical interaction with the sample, multi-target imaging was realized with light-induced activation of fluorescent probes, e.g. using photoactivatable or photoconvertible fluorescent proteins^[14– 16]^ and fluorophore-specific activation with light.^[17]^ Photoactivatable synthetic organic dyes were also engineered for single-molecule localization microscopy and STED microscopy.^[18–22]^ However, in all these approaches, fluorophores are covalently attached to their target structures, and selective removal of a signal requires photobleaching.

Here, we bridge the gap between the exchangeable nature of DNA-based protein labeling and using light as a non-invasive tool to control molecular targets in biological systems.^[23,24]^ Unlike previous methods that modulate fluorophore photophysics, we present a strategy that uses photo-modulatable groups (photocages) integrated into short DNA sequences to control the hybridization state with light. Such photocages have previously been used in the DNA hairpin structures to modulate the cellular activity of short DNA oligonucleotides.^[25,26]^ Here, we report the integration of photocages in short DNA oligonucleotides and demonstrate wash-free switching between imaging targets using light. We demonstrate this concept for confocal microscopy, single-molecule localization microscopy (SMLM), and stimulated emission depletion (STED) microscopy.

## Results and Discussion

The coumarin photocage derivative methyl-7-(diethylamino)coumarin-4-ylmethyl (Me-DEACM) was chosen for the construction of photo-modulatable DNA oligonucleotides. Coumarin photocages release their payload by irradiation with visible light and in a well-behaved manner, minimizing cell damage.^[27]^ For application in oligonucleotides, commercially available phosphoramidites are available. Thus, coumarin photocages are commonly applied and have previously been used masking the function of oligonucleotides or amino acids e.g., to modulate mRNA translation,^[28]^ monitor mRNA transport in neurons,^[29]^ or to gain insights into zebrafish embryo development by light-regulation of proteins.^[30]^ The absorption maximum in the blue-violet spectral region is orthogonal to commonly employed orange- and red-absorbing fluorophores in fluorescence microscopy.

In this work, we selected a methylated DEACM photocage, as it provides a slightly higher uncaging quantum yield compared to the unmodified DEACM.^[31]^ We opted for two orthogonal, photocaged oligonucleotide docking strands that simultaneously alter their hybridization properties upon irradiation (**Figure 1A**). Docking strand **cP1**_**a**_ is in an active state (“ON” ) before uncaging and allows hybridization with the imager strand P1 and imaging of the first target. At the same time, the imager strand cannot bind to docking strand **cP1**_**b**_ as the interaction is sterically hindered by the coumarin groups (“OFF” ). After irradiation with violet light, the system is reversed: the photocages are released, and **cP1**_**a**_ internally hybridizes, yielding a closed, inactive docking strand (“OFF” ), while **cP1**_**b**_ adopts an active conformation, enabling imaging of the second target (“ON” ).

**Figure 1.**
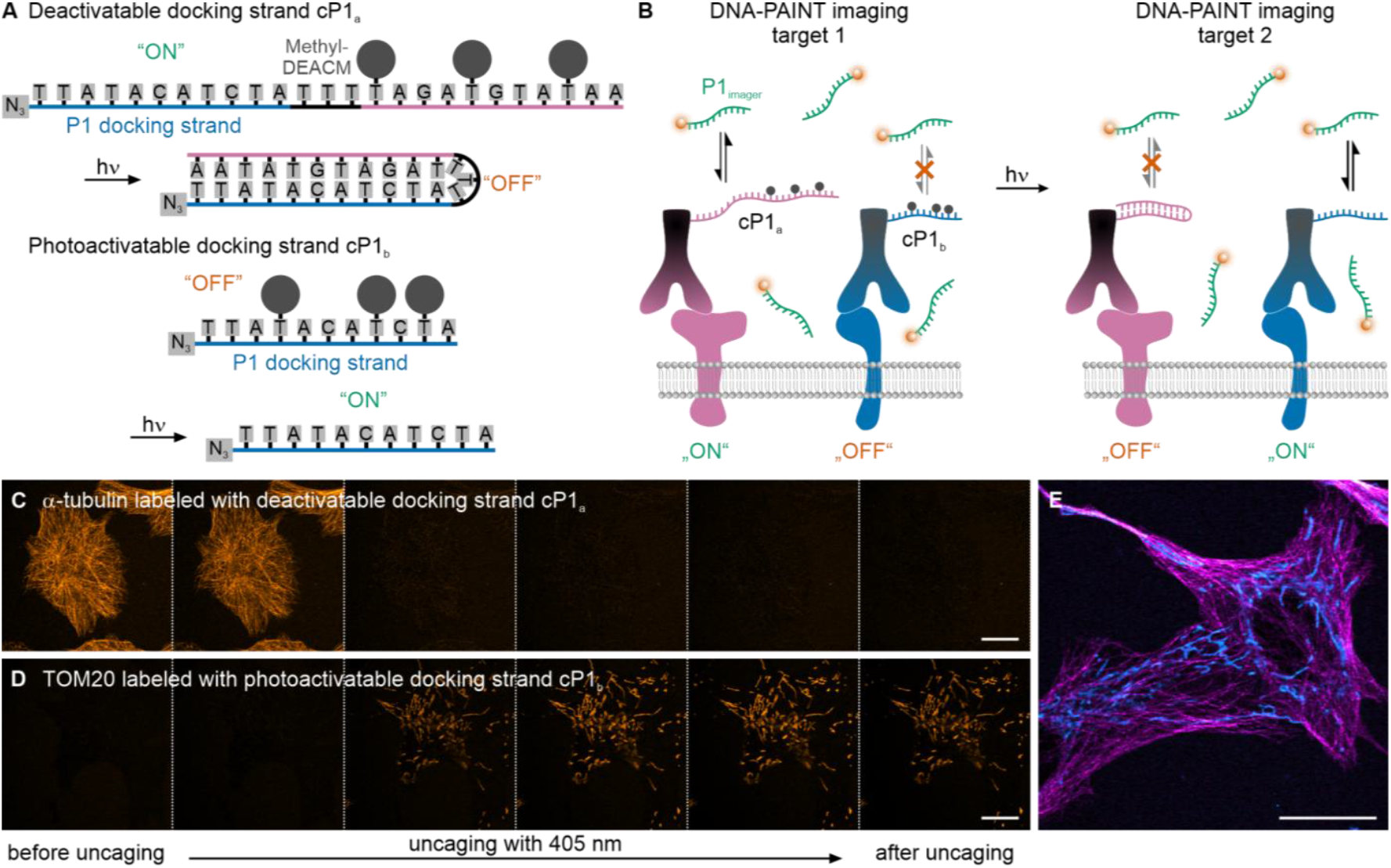
Wash-free DNA-PAINT using caged docking strands. (A) Design of the photodeactivatable and photoactivatable docking strands **cP1**_**a**_ and **cP1**_**b**_. Initially, **cP1**_**a**_ is accessible for hybridization to the complementary P1 imager strand, while **cP1**_**b**_ remains in its “OFF” state. Upon 405-nm illumination, the Methyl-DEACM cage groups are removed: **cP1**_**a**_ adopts a closed, inactive (“OFF” ) conformation, whereas **cP1**_**b**_ becomes activated and available for P1 imager binding. (B) Scheme of the wash-free two-target imaging workflow. Each target is labeled with either **cP1**_**a**_ or **cP1**_**b**_, and images are recorded before and after 405-nm illumination. (C) Confocal images of α-tubulin labeled with **cP1**_**a**_ show loss of microtubule signal during illumination with violet light. (D) Confocal images of TOM20 labeled with **cP1**_**b**_ show the appearance of the mitochondrial signal following illumination. All images were acquired using 100 nM P1-Cy3B imager strand. (E) Two-target confocal imaging of microtubules (magenta) and mitochondria (cyan). α-Tubulin was labeled with **cP1**_**a**_, and TOM20 with **cP1**_**b**_. The two-target image was generated by combining images recorded before and after violet-light illumination. Imager strand concentration was 100 nM P1-Cy3B. Scale bars 20 µm.

### Synthesis

The chemical synthesis (**Scheme 1**) started with a Grignard reaction of **1**, affording the methylated DEACM **2**. Next, **2** was reacted with the activated dT derivative **S3** (**Scheme S1**) to attach the coumarin moiety to the O4 and hence inhibit H-bonds between the nucleobases. Here, it is important to verify that the coumarin is connected to O4 using NMR, as in the case of attachment to N3, the rate of photolysis is strongly decreased. With the right isomer in hand, the tert-butyldimethylsilyl (TBDMS) protecting groups of **3** were cleaved, followed by attachment of dimethoxytrityl (DMTr) protecting group to the 5’ position and finally phosphitylation to obtain the caged thymidine **4**. A comprehensive description of the synthesis is found in the **Supporting Information**. Compound **4** was then used in the solid-phase synthesis of the oligonucleotides. Both oligonucleotides were designed to contain the widely used P1-docking strand sequence.^[5,32–34]^ For **cP1**_**b**,_ three positions were chosen to introduce caged dT **4** and consequently disable hybridization with an imager strand (**Figure 1A**). For **cP1**_**a**_, the P1 docking strand was extended with a fully complementary sequence containing three caged dTs to ensure complete internal hybridization after irradiation. Post-synthetically, an azide modification was introduced for antibody labelling.

**Scheme 1.**
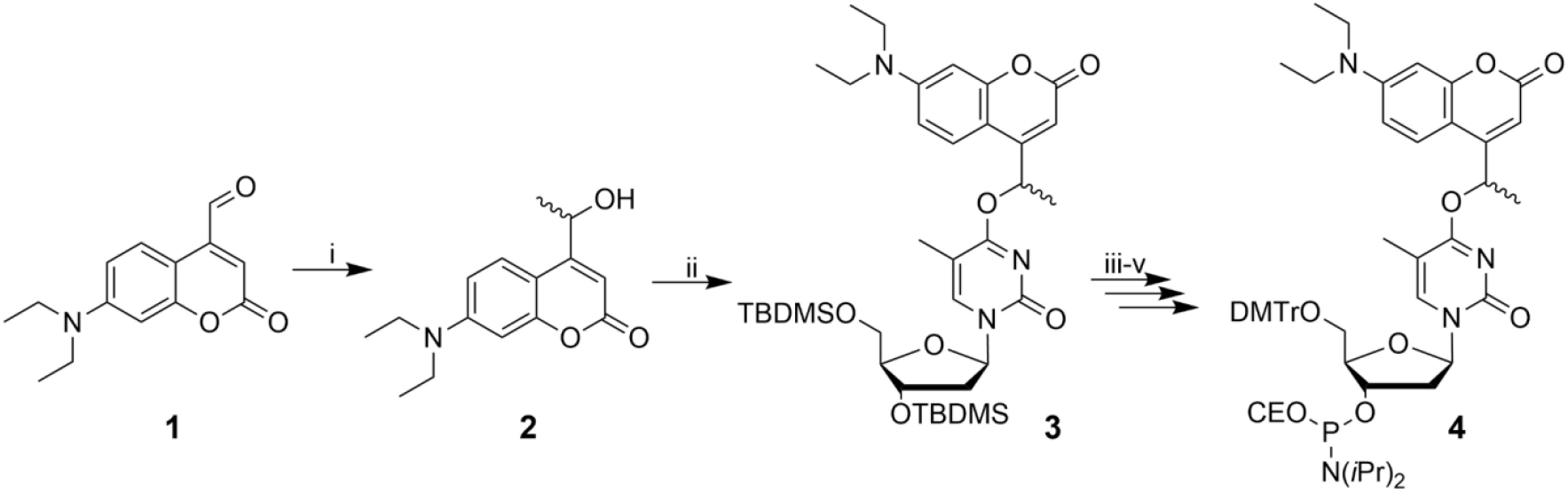
Synthesis of Methyl-DEACM caged dT-Phosphoramidite **4**. i. MeMgBr, THF, -78 °C to rt, 21 h, 41%. ii. **S3**, DBU, ACN, rt, 20 h, 77%. iii. TBAF, AcOH, THF, 0 °C to rt, 20 h, 85%. iv. DMTr-Cl, DIPEA, DCM, 0 °C to rt, on, 83%. v. PN(*i*Pr)_2_(CEO)-Cl, DIPEA, DCM, 0 °C to rt, 1.5 h, 80%.

We performed *in vitro* photolysis experiments with the caged docking strands **cP1**_**a**_ and **cP1**_**b**_ (**Figure S1 and S2**) *via* RP-HPLC and uridine as an internal standard. Both caged docking strands showed rapid photolysis already after irradiation for 15 s at 405 nm (P = 19.8 mW), and signals of the docking strand and the coumarin photocage appeared. After completing irradiation, the photolysis products were analyzed and quantified using the internal standard. For **cP1**_**a**_ 50% and for **cP1**_**b**_ 71% of the uncaged docking strand were present. Both photolysis indicated two coumarin signals, which we assume to be Methyl-DEACM-OH and a red-shifted elimination derivative according to previous literature.^[31]^

### Caged Docking Strands for Wash-Free Multiplexed Fluorescence Microscopy

The caged docking strands **cP1**_**a**_ and **cP1**_**b**_ (**Figure 1A**) were conjugated to secondary antibodies via click chemistry (see **Experimental Procedures**) and used to label two distinct cellular structures by immunofluorescence. The first target was immunolabeled with **cP1**_**a**_ and imaged by transiently hybridizing to a fluorophore-labeled P1 imager strand. To switch from the first to the second target, low-intensity illumination with violet light (405 nm) was applied, which removes the Methyl-DEACM caging groups from **cP1**_**a**_ and **cP1**_**b**_. This leads to an intramolecular hybridization of **cP1**_**a**_, rendering it inaccessible for imaging, and makes **cP1**_**b**_ accessible for imaging the second target with the same fluorophore-labeled P1 imager strand (**Figure 1B**). In contrast to Exchange-DNA-PAINT,^[5]^ no exchange of the imager strand is necessary for imaging the second target.

To validate the functionality of the caged docking strands, docking-strand–conjugated antibodies were used to label individual targets in U-2 OS cells (**Figure 1C,D**). α-tubulin was labeled with the deactivatable docking strand **cP1**_**a**_, and confocal imaging showed the expected microtubule network before uncaging. Upon illumination with very low intensity (radiant exposure: ∼0.3-0.6 J/cm^2^) of 405-nm light, the tubulin signal disappeared, demonstrating highly efficient uncaging. TOM20 was labeled using the photoactivatable docking strand **cP1**_**b**_, and the mitochondrial network became visible only after uncaging. Again, only a few frames of illumination with 405 nm were sufficient to reveal the mitochondrial structure. To quantify the uncaging efficiency in cells, we analyzed confocal images of α-tubulin labeled with **cP1**_**a**_ and **cP1**_**b**_ before, during, and after violet-light illumination. After illumination for 5 frames with 405-nm light, the signal disappeared for **cP1**_a_ and saturated for **cP1**_**b**_ (**Figure S3**).

After confirming that both docking strands were efficiently switched between their “ON” and “OFF” states, we labeled microtubules and mitochondria within the same sample and demonstrated that both structures can be visualized (**Figure 1E**). During uncaging, the transition from the microtubule to mitochondrial signal was clearly observable (**Figure S4A** and **Video S1**). The **cP1**_**a**_**/cP1**_**b**_ system was subsequently applied to additional cellular targets (**Figure S4B–D**). In comparison to conventional DNA-PAINT experiments, this approach avoids washing steps (typically 3–5 phosphate-buffered saline washes with waiting times of 5–10 min in between) as well as the addition of new imager strands. As a result, two-target imaging using fluorophore-labeled oligonucleotides can be performed without physically perturbing the nanoscale topography of the sample. In addition, using light to switch between targets enables spatially precise conversion (**Figure S5**).

Next, we tested the compatibility of the caged docking strands with additional cellular stains across the visible spectrum. Microtubules and vimentin were labeled using **cP1**_**a**_ and **cP1**_**b**_, respectively, using an ATTO 655-labeled P1 imager strand. Additionally, actin was stained with ATTO 488-phalloidin and mitochondria with MitoTracker Orange. This configuration enabled the imaging of four distinct targets using only three spectral channels (**Figure 2A**), demonstrating that caged docking strands can expand the number of addressable targets in otherwise spectrally limited experiments.

**Figure 2.**
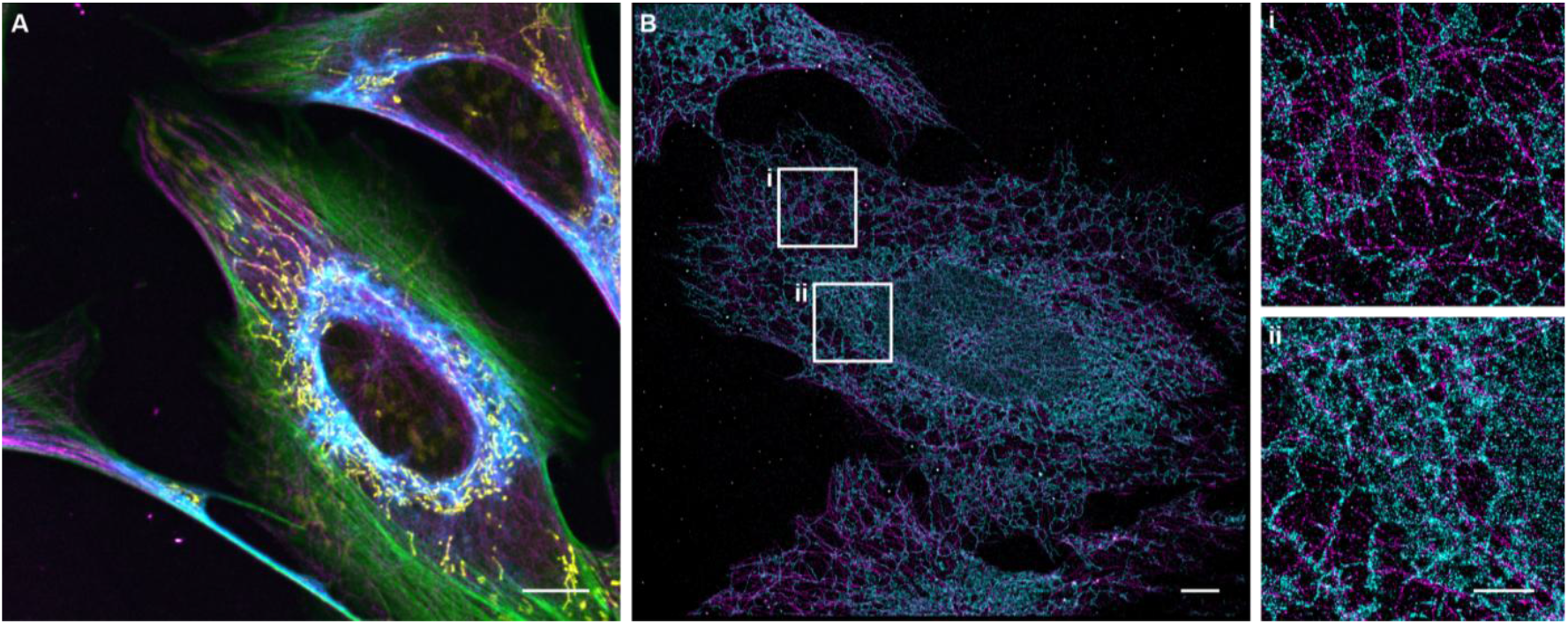
(A) Multiplexed confocal imaging of microtubules, vimentin, actin, and mitochondria in U-2 OS cells. α-tubulin (magenta) and vimentin (cyan) were labeled via immunofluorescence using **cP1**_**a**_ and **cP1**_**b**_ conjugated secondary antibodies, respectively, in combination with 50 nM P1(11nt)-ATTO 655. Actin was stained with ATTO 488-Phalloidin (green) and mitochondria with MitoTracker Orange (yellow). Scale bar 10 µm. (B) Two-target DNA-PAINT imaging of microtubules and ER in U-2 OS cells with caged docking strands. α-tubulin was labeled with **cP1**_**a**_ (magenta), and KDEL was labeled with **cP1**_**b**_ (cyan). 2 nM P1(9nt)-Cy3B imager strand was used for imaging. Before imaging the second target, the cell was illuminated for 2 min with 405-nm light (radiant exposure: ∼600 J/cm^2^). The NeNA (nearest-neighbor analysis) localization precision for the DNA-PAINT image of α-tubulin was 5.33 nm, and for KDEL 5.47 nm. Scale bar 5 µm, zoom-ins 2 µm.

To show the applicability of caged docking strands to super-resolution microscopy, two-target DNA-PAINT experiments were performed (**Figure 2B, Figure S6**). Microtubules and ER were labeled with **cP1**_**a**_ and **cP1**_**b**_, respectively. First, α-tubulin was imaged using a P1-Cy3B imager strand. After recording the first DNA-PAINT movie, the photocages were removed by illuminating the cell for 2 min with 405-nm light. Next, the KDEL sequence residing in the ER was imaged using the same imager strand. The two-target super-resolution image shows the nanoscale co-organization of microtubules and ER (**Figure 2Bi,ii**), in line with previous reports that the ER interacts with microtubules, sliding along stable, acetylated microtubules to help regulate ER positioning and organelle distribution.^[35,36]^

DNA-fluorophore labels were previously used for STED microscopy.^[10,37]^ We found that caged docking strands are equally applicable, and demonstrated two-target STED imaging using **cP1**_**a**_ and **cP1**_**b**_ (**Figure S7**). To increase the signal-to-background ratio, we used self-quenched imager strands dual-labeled with Cy3B, which have a lower background fluorescence signal in the unbound state and show an increase in fluorescence signal upon binding to the complementary docking strand.^[38,39]^ Hence, two-target STED microscopy using a single type of imager strand was possible (**Figure S7**), which adds a new promising strategy for multi-target super-resolution microscopy. We observed that the 775-nm depletion laser caused partial uncaging of Methyl-DEACM, which needs to be considered when using caged docking strands for STED microscopy (**Supporting Note 1**).

## Conclusion

We introduce cagePAINT as a strategy for wash-free, multi-target high-resolution imaging. By integrating photocages into DNA oligonucleotides, we demonstrate two-target confocal, DNA-PAINT, and STED microscopy using a single fluorophore-labeled imager strand. The highly efficient uncaging process triggered by violet light allows locally confined conversion, enabling dual-target imaging on a cell-by-cell basis and accelerating DNA-PAINT acquisition by eliminating the need for washing steps between imaging rounds. The number of targets can be increased by using photocages of different wavelengths, profiting e.g. from highly selective uncaging of coumarin-based caging groups with minimal cross-activation.^[40]^ Additionally, a variety of photocages is already established,^[27]^ including red-light activatable ones like BODIPY, cyanines, or xanthenes.^[41]^ At the same time, orthogonality includes more than a minimal cross-section,^[42]^ and other possibilities are simultaneous or consecutive two-photon excitation.^[43,44]^ Caged oligonucleotides can also be integrated into existing methods for DNA-based multi-target microscopy and further increase multiplexing.^[12,13]^ The concept of photocages for multi-target imaging is not limited to DNA-fluorophore labels, and can be extended to photocaged self-labelling enzymes.^[45]^ The integration into other microscopy modalities, such as structured-illumination microscopy (SIM) or MINFLUX, is possible.^[46]^ In summary, cagePAINT is economical, as only a single type of imager strand is used, and eliminates physical distortions and sample drift that can occur during washing in multiplexed DNA-PAINT experiments. Consequently, the accuracy and thus information content of super-resolution structural data are enhanced.

## Supporting Information

The authors have cited additional references within the Supporting Information.^[47–51]^

## Acknowledgements

We thank Petra Freund for assistance with cell culture, Tanja Menche for helpful discussions, Jonathan Marheineke for assistance in chemical synthesis, and Till Stephan for support with STED microscopy. We gratefully acknowledge funding by the Deutsche Forschungsgemeinschaft (DFG, German Research Foundation) through CRC 1507, DFG INST 161/926-1 FUGG, DFG INST 161/778-1 FUGG, and DFG INST 161/1162-1 FUGG.

## Notes

### Competing Interest Statement

The authors have declared no competing interest.

### Summary of Updates

Figure 1 revised, new Figure S3 and S5

## References

[1] J. S. H. Danial, Nat Methods 2025, 22, 1636–1652.

[2] H. G. Schroeter, S. Sass, M. Heilemann, T. Kuner, M. Klevanski, Adv Sci (Weinh) 2025, e06731.

[3] M. Klevanski, F. Herrmannsdoerfer, S. Sass, V. Venkataramani, M. Heilemann, T. Kuner, Nat Commun 2020, 11, 1552.

[4] L. Albertazzi, M. Heilemann, Angew Chem Int Ed Engl 2023, 62, e202303390.

[5] R. Jungmann, M. S. Avendaño, J. B. Woehrstein, M. Dai, W. M. Shih, P. Yin, Nat Methods 2014, 11, 313–318.

[6] J. Kompa, J. Bruins, M. Glogger, J. Wilhelm, M. S. Frei, M. Tarnawski, E. D’Este, M. Heilemann, J. Hiblot, K. Johnsson, J Am Chem Soc 2023, 145, 3075–3083.

[7] J. Kompa, L. J. Dornfeld, N. Porzberg, S. Jang, S. H. Lilje, C. Catapano, D. Jocher, L. Merk, S. Zedlitz, R. Mao, J. Wilhelm, M. Dietz, M. Tarnawski, J. Hiblot, M. Heilemann, K. Johnsson, bioRxiv 2025, DOI 10.1101/2025.07.06.663254.

[8] C. K. Spahn, M. Glaesmann, J. B. Grimm, A. X. Ayala, L. D. Lavis, M. Heilemann, Sci Rep 2018, 8, 14768.

[9] M. Glogger, D. Wang, J. Kompa, A. Balakrishnan, J. Hiblot, H.-D. Barth, K. Johnsson, M. Heilemann, ACS Nano 2022, 16, 17991–17997.

[10] C. Spahn, F. Hurter, M. Glaesmann, C. Karathanasis, M. Lampe, M. Heilemann, Angew Chem Int Ed Engl 2019, 58, 18835–18838.

[11] M. Glogger, C. Spahn, J. Enderlein, M. Heilemann, Angew Chem Int Ed Engl 2021, 60, 6310–6313.

[12] E. M. Unterauer, S. Shetab Boushehri, K. Jevdokimenko, L. A. Masullo, M. Ganji, S. Sograte-Idrissi, R. Kowalewski, S. Strauss, S. C. M. Reinhardt, A. Perovic, C. Marr, F. Opazo, E. F. Fornasiero, R. Jungmann, Cell 2024, 187, 1785–1800.e16.

[13] F. Schueder, F. Rivera-Molina, M. Su, Z. Marin, P. Kidd, J. E. Rothman, D. Toomre, J. Bewersdorf, Cell 2024, 187, 1769–1784.e18.

[14] G. H. Patterson, J. Lippincott-Schwartz, Science 2002, 297, 1873–1877.

[15] F. V. Subach, G. H. Patterson, S. Manley, J. M. Gillette, J. Lippincott-Schwartz, V. V. Verkhusha, Nature Methods 2009, 6, 153–159.

[16] D. M. Shcherbakova, P. Sengupta, J. Lippincott-Schwartz, V. V. Verkhusha, Annu Rev Biophys 2014, 43, 303–329.

[17] D. Virant, B. Turkowyd, A. Balinovic, U. Endesfelder, Int J Mol Sci 2017, 18, DOI 10.3390/ijms18071524.

[18] J. B. Grimm, B. P. English, H. Choi, A. K. Muthusamy, B. P. Mehl, P. Dong, T. A. Brown, J. Lippincott-Schwartz, Z. Liu, T. Lionnet, L. D. Lavis, Nature Methods 2016, 13, 985–988.

[19] M. S. Frei, P. Hoess, M. Lampe, B. Nijmeijer, M. Kueblbeck, J. Ellenberg, H. Wadepohl, J. Ries, S. Pitsch, L. Reymond, K. Johnsson, Nature Communications 2019, 10, 4580.

[20] M. Weber, T. A. Khan, L. J. Patalag, M. Bossi, M. Leutenegger, V. N. Belov, S. W. Hell, Chemistry 2021, 27, 451–458.

[21] R. Lincoln, M. L. Bossi, M. Remmel, E. D’Este, A. N. Butkevich, S. W. Hell, Nat Chem 2022, 14, 1013–1020.

[22] I. Likhotkin, R. Lincoln, M. L. Bossi, A. N. Butkevich, S. W. Hell, J Am Chem Soc 2023, 145, 1530–1534.

[23] C. Brieke, F. Rohrbach, A. Gottschalk, G. Mayer, A. Heckel, Angew. Chem. Int. Ed Engl. 2012, 51, 8446–8476.

[24] A. E. Mangubat-Medina, Z. T. Ball, Chem. Soc. Rev. 2021, 50, 10403–10421.

[25] S. Sambandan, G. Akbalik, L. Kochen, J. Rinne, J. Kahlstatt, C. Glock, G. Tushev, B. Alvarez-Castelao, A. Heckel, E. M. Schuman, Science 2017, 355, 634–637.

[26] T. Lucas, F. Schäfer, P. Müller, S. A. Eming, A. Heckel, S. Dimmeler, Nat Commun 2017, 8, 15162.

[27] R. Weinstain, T. Slanina, D. Kand, P. Klán, Chem. Rev. 2020, 120, 13135–13272.

[28] A. Bollu, N. Klöcker, P. Špaček, F. P Weissenboeck, S. Hüwel, A. Rentmeister, Angew. Chem. Int. Ed Engl. 2023, 62, e202209975.

[29] R. Klimek, P. G. Donlin-Asp, C. Polisseni, V. Hanff, E. M. Schuman, A. Heckel, Chem. Commun. (Camb.) 2021, 57, 12683–12686.

[30] W. Brown, J. Wesalo, S. Samanta, J. Luo, S. E. Caldwell, M. Tsang, A. Deiters, ACS Chem. Biol. 2023, 18, 1305–1314.

[31] A. M. Schulte, G. Alachouzos, W. Szymański, B. L. Feringa, J. Am. Chem. Soc. 2022, 144, 12421–12430.

[32] J. Schnitzbauer, M. T. Strauss, T. Schlichthaerle, F. Schueder, R. Jungmann, Nat Protoc 2017, 12, 1198–1228.

[33] K. K. Narayanasamy, J. V. Rahm, S. Tourani, M. Heilemann, Nat Commun 2022, 13, 5047.

[34] N. Oleksiievets, Y. Sargsyan, J. C. Thiele, N. Mougios, S. Sograte-Idrissi, O. Nevskyi, I. Gregor, F. Opazo, S. Thoms, J. Enderlein, R. Tsukanov, Communications Biology 2022, 5, 38.

[35] J. R. Friedman, B. M. Webster, D. N. Mastronarde, K. J. Verhey, G. K. Voeltz, J Cell Biol 2010, 190, 363–375.

[36] P. Zheng, C. J. Obara, E. Szczesna, J. Nixon-Abell, K. K. Mahalingan, A. Roll-Mecak, J. Lippincott-Schwartz, C. Blackstone, Nature 2021, 601, 132–138.

[37] S. Beater, P. Holzmeister, B. Lalkens, P. Tinnefeld, Opt Express 2015, 23, 8630–8638.

[38] L. F. Kessler, A. Balakrishnan, T. Menche, D. Wang, Y. Li, M. Mantel, M. Glogger, M. S. Dietz, M. Heilemann, J Phys Chem B 2024, 128, 6751–6759.

[39] L. F. Kessler, A. Balakrishnan, N. S. Deußner-Helfmann, Y. Li, M. Mantel, M. Glogger, H.-D. Barth, M. S. Dietz, M. Heilemann, Angew Chem Int Ed Engl 2023, 62, e202307538.

[40] J. Kaufmann, F. Sinsel, A. Heckel, Chemistry 2023, 29, e202204014.

[41] A. Egyed, K. Németh, T. Á. Molnár, M. Kállay, P. Kele, M. Bojtár, J Am Chem Soc 2023, 145, 4026–4034.

[42] F. Pashley-Johnson, X. Wu, J. A. Carroll, S. L. Walden, H. Frisch, A.-N. Unterreiner, F. E. Du Prez, H.-A. Wagenknecht, J. Read de Alaniz, B. L. Feringa, A. Heckel, C. Barner-Kowollik, Angew Chem Int Ed Engl 2025, 64, e202502651.

[43] Y. Becker, E. Unger, M. A. H. Fichte, D. A. Gacek, A. Dreuw, J. Wachtveitl, P. J. Walla, A. Heckel, Chem. Sci. 2018, 9, 2797–2802.

[44] L. J. G. W. van Wilderen, D. Kern-Michler, C. Neumann, M. Reinfelds, J. von Cosel, M. Horz, I. Burghardt, A. Heckel, J. Bredenbeck, Chem Sci 2023, 14, 2624–2630.

[45] F. Walterspiel, B. Ugarte-Uribe, J. Weidenhausen, M. Vincent, K. K. Narayanasamy, A. Dimitriadi, A. U. M. Khan, M. Fritsch, C. W. Müller, T. Zimmermann, C. Deo, Angew Chem Int Ed Engl 2025, e202424955.

[46] N. Radmacher, A. I. Chizhik, O. Nevskyi, J. I. Gallea, I. Gregor, J. Enderlein, Annu Rev Biophys 2025, 54, 163–184.

[47] P. Seyfried, L. Eiden, N. Grebenovsky, G. Mayer, A. Heckel, Angew Chem Int Ed Engl 2017, 56, 359–363.

[48] A. D. Edelstein, M. A. Tsuchida, N. Amodaj, H. Pinkard, R. D. Vale, N. Stuurman, J Biol Methods 2014, 1, DOI 10.14440/jbm.2014.36.

[49] U. Endesfelder, S. Malkusch, F. Fricke, M. Heilemann, Histochem Cell Biol 2014, 141, 629–638.

[50] J. Vogelsang, R. Kasper, C. Steinhauer, B. Person, M. Heilemann, M. Sauer, P. Tinnefeld, Angew Chem Int Ed Engl 2008, 47, 5465–5469.

[51] P. Blumhardt, J. Stein, J. Mücksch, F. Stehr, J. Bauer, R. Jungmann, P. Schwille, Molecules 2018, 23, DOI 10.3390/molecules23123165.

