## Supplemental Information for "Wash-Free Multi-Target Super-Resolution Microscopy with Photocaged DNA Labels"

#These authors contributed equally.

#### Table of Contents

**Supporting Note** **2**
**Supporting Figures** **3**
**Supporting Video** **8**
**Experimental Procedures** **9**
    Materials and Methods 9
    Chemical Synthesis 10
    Oligonucleotide Synthesis 14
    Sample Preparation for Antibody Labeling 15
    Photochemical Characterization 17
    Labeling of Antibodies with Photocaged Docking Strands 17
    Confocal Laser Scanning Microscopy 19
    DNA-PAINT Imaging 19
    STED Microscopy 20
**Appendix** **21**
    NMR Spectra 21
    Mass Spectra 27
    Oligonucleotide Mass Spectra 29
**References** **31**

### Supporting Note

#### Supporting Note 1

We performed STED microscopy using an excitation laser at 561 nm and a depletion laser at 775 nm. Using these imaging settings, we observed that high intensities of the STED depletion laser (775 nm) can induce partial uncaging of Methyl-DEACM. At the depletion laser intensity used in this work (~90 MW/cm²), partial uncaging was detected after the measurement. However, no effect was apparent during image acquisition (**Figure S7**). At lower depletion laser intensities (~35 MW/cm²), no uncaging was observed during the measurement of the first target. We therefore recommend careful control and adjustment of the imaging settings to prevent partial uncaging during imaging of the first target.

### Supporting Figures

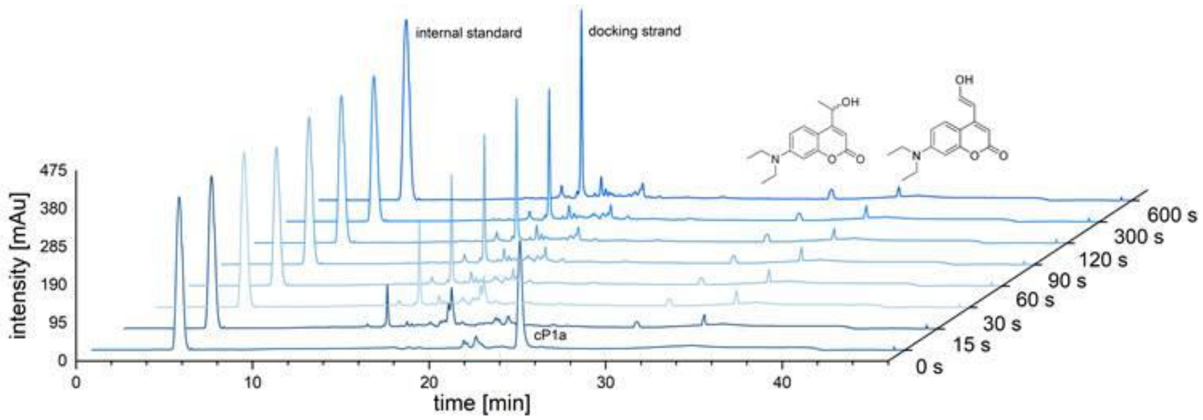


**Figure S1.** RP-HPLC chromatograms (254 nm traces) of photolysis of **cP1a**. Irradiation=405 nm (P=19.8 mW), c=2.5 µM, n=2 nmol, 1x PBS (pH 7.4), internal standard=uridine (c=100 µM).

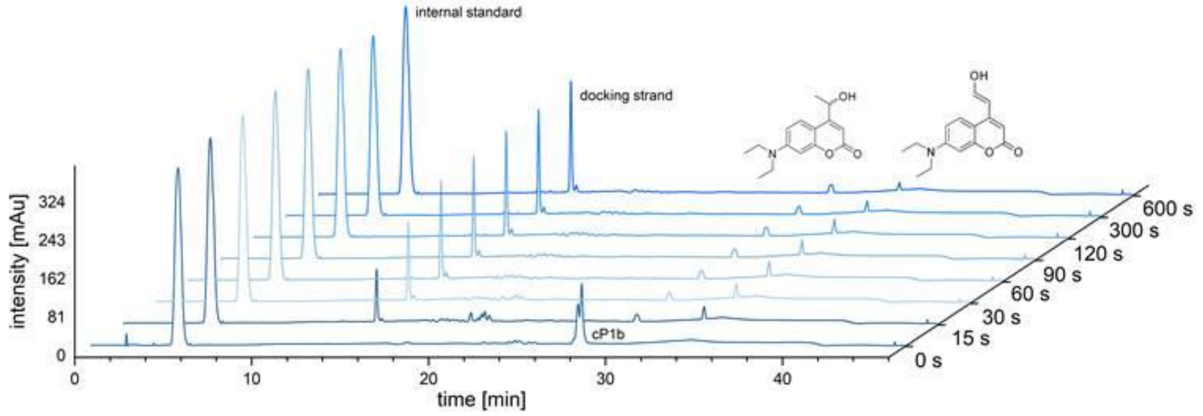


**Figure S2.** RP-HPLC chromatograms (254 nm traces) of photolysis of **cP1b**. Irradiation=405 nm (P=19.8 mW), c=2.5 µM, n=2 nmol, 1x PBS (pH 7.4), internal standard=uridine (c=100 µM).

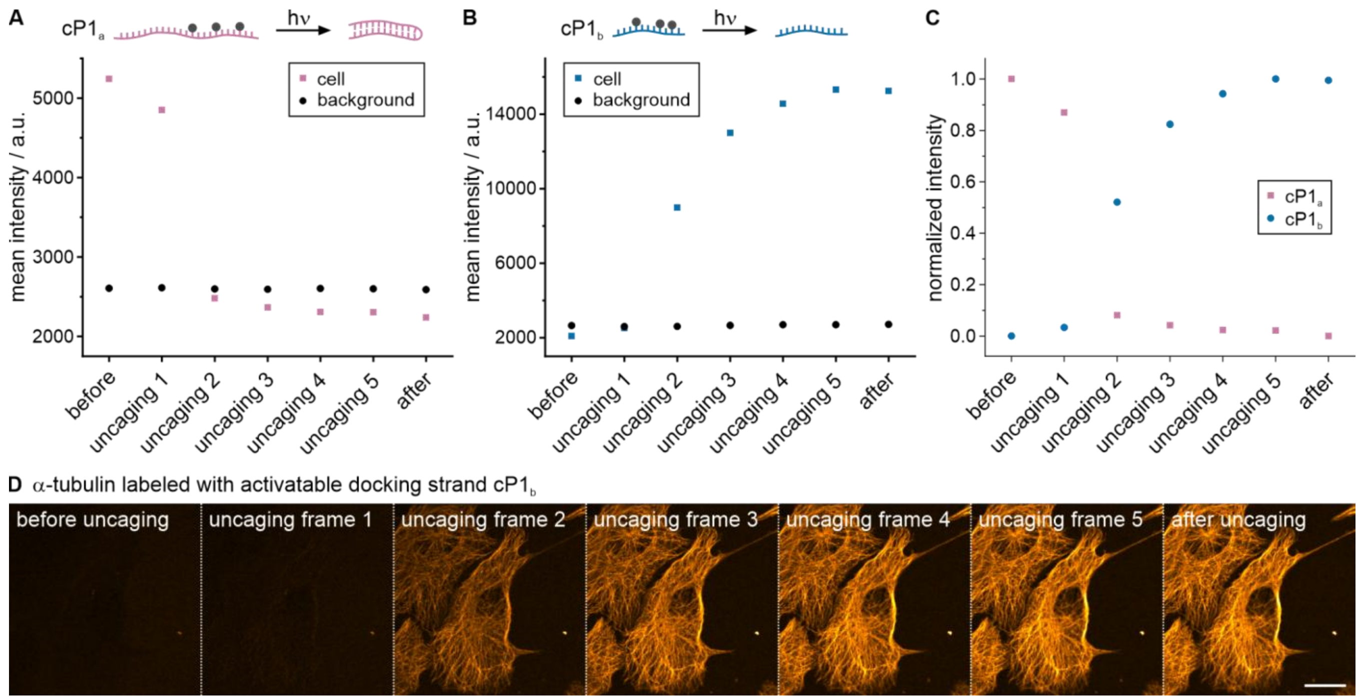


**Figure S3.** Quantitative analysis of the uncaging process of **cP1a** and **cP1b** oligonucleotides extracted from confocal laser scanning microscopy data. U-2 OS cells were immunostained for α-tubulin using antibodies conjugated to **cP1a** and **cP1b** oligonucleotides and fluorescently labeled with the P1-Cy3B imager strand (100 nM). The fluorescence intensity before, during, and after illumination with 405-nm light was quantified frame-wise by drawing a region of interest around a target cell and a background region. (A) The fluorescence intensity decreases upon uncaging of **cP1a**. The cell shown in **Figure 1C** was analyzed. (B) The fluorescence intensity increases upon uncaging of **cP1b**. The cell shown in **Figure S3D** was analyzed. (C) Comparison of the normalized intensity change of **cP1a** and **cP1b**. (D) Confocal images of α-tubulin labeled with **cP1b** show an increase in microtubule signal upon illumination with violet light. Scale bar 20 µm.

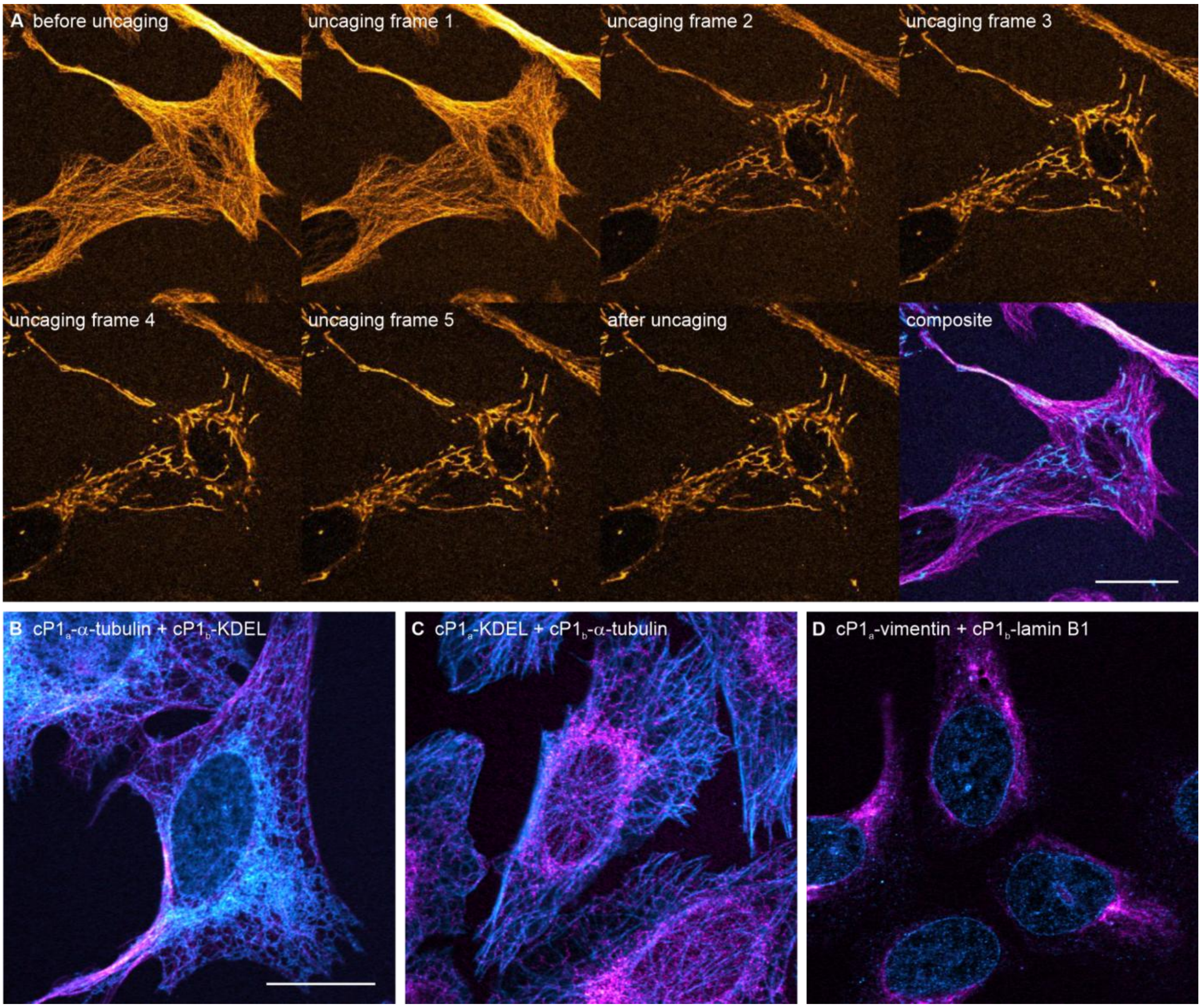


**Figure S4.** Two-target confocal imaging with caged docking strands. (A) Time series of α-tubulin labeled with $\mathbf{cP1_a}$ and TOM20 labeled with $\mathbf{cP1_b}$. The confocal image before uncaging, the uncaging process upon illumination with 405 nm, the confocal image after uncaging, and the composite two-target image (α-tubulin: magenta, TOM20: cyan) are shown. The composite image was generated from images acquired before and after violet-light illumination. The uncaging process is highly efficient, such that the second structure is already visible in the second frame of 405 nm illumination. (B-D) Two-target confocal images of α-tubulin labeled with $\mathbf{cP1_a}$ (magenta) and KDEL labeled with $\mathbf{cP1_b}$ (cyan), KDEL labeled with $\mathbf{cP1_a}$ (magenta) and α-tubulin labeled with $\mathbf{cP1_b}$ (cyan), and vimentin labeled with $\mathbf{cP1_a}$ (magenta) and lamin B1 labeled with $\mathbf{cP1_b}$ (cyan). All images were acquired with 100 nM P1-Cy3B imager strand. Scale bars 20 µm.

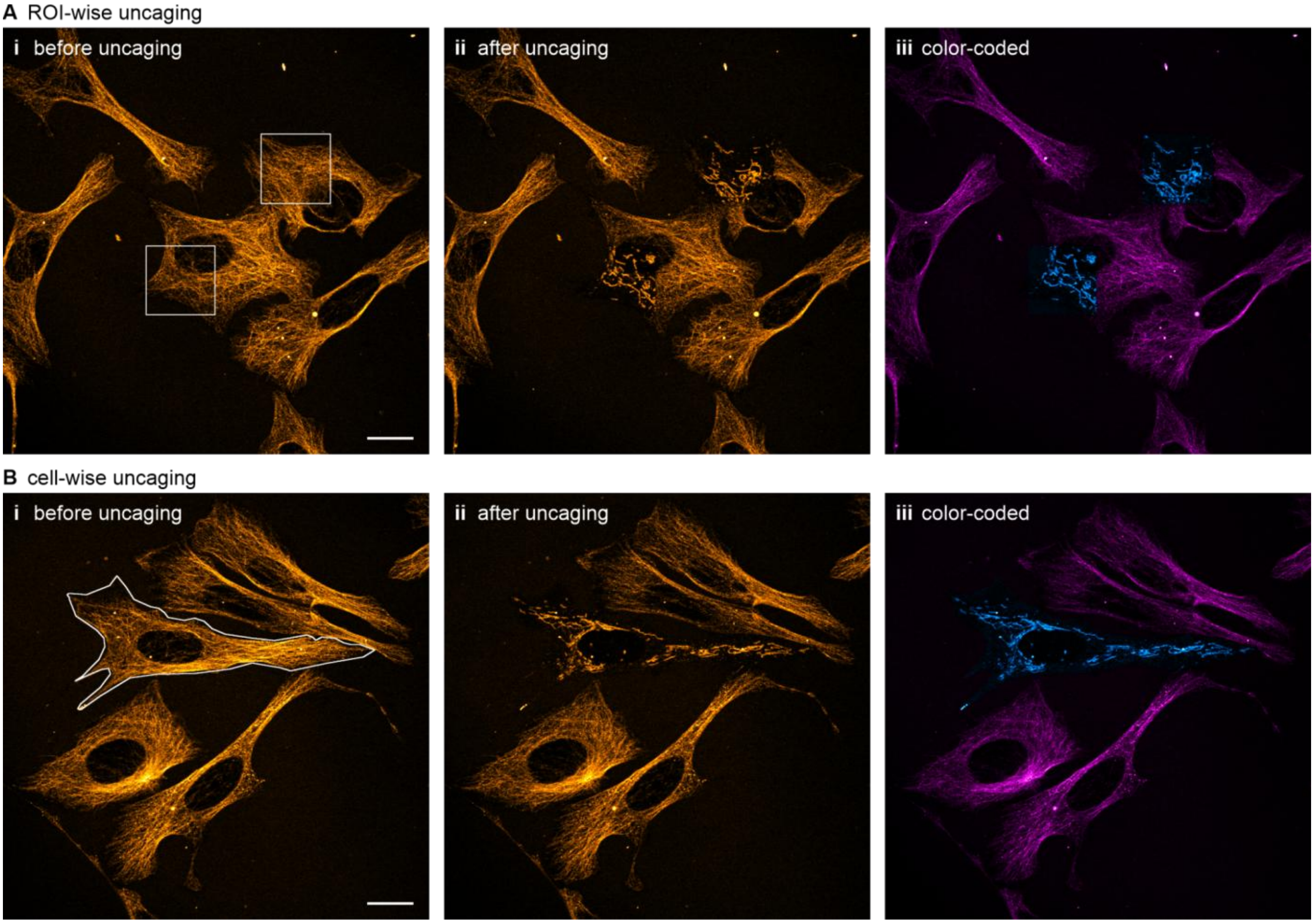


**Figure S5.** Spatially confined two-target confocal imaging using caged docking strands. α-Tubulin and TOM20 were labeled with **cP1a** and **cP1b**, respectively, and 100 nM of P1-ATTO643 was added to the imaging buffer. Spatially selective uncaging is shown (A) for subcellular regions or (B) entire individual cells. Confocal images acquired before uncaging (i) and after 405-nm illumination and in the highlighted ROI (ii) are shown, together with a color-coded image generated from both recordings (α-tubulin: magenta, TOM20: cyan) (iii). Scale bars 20 μm.

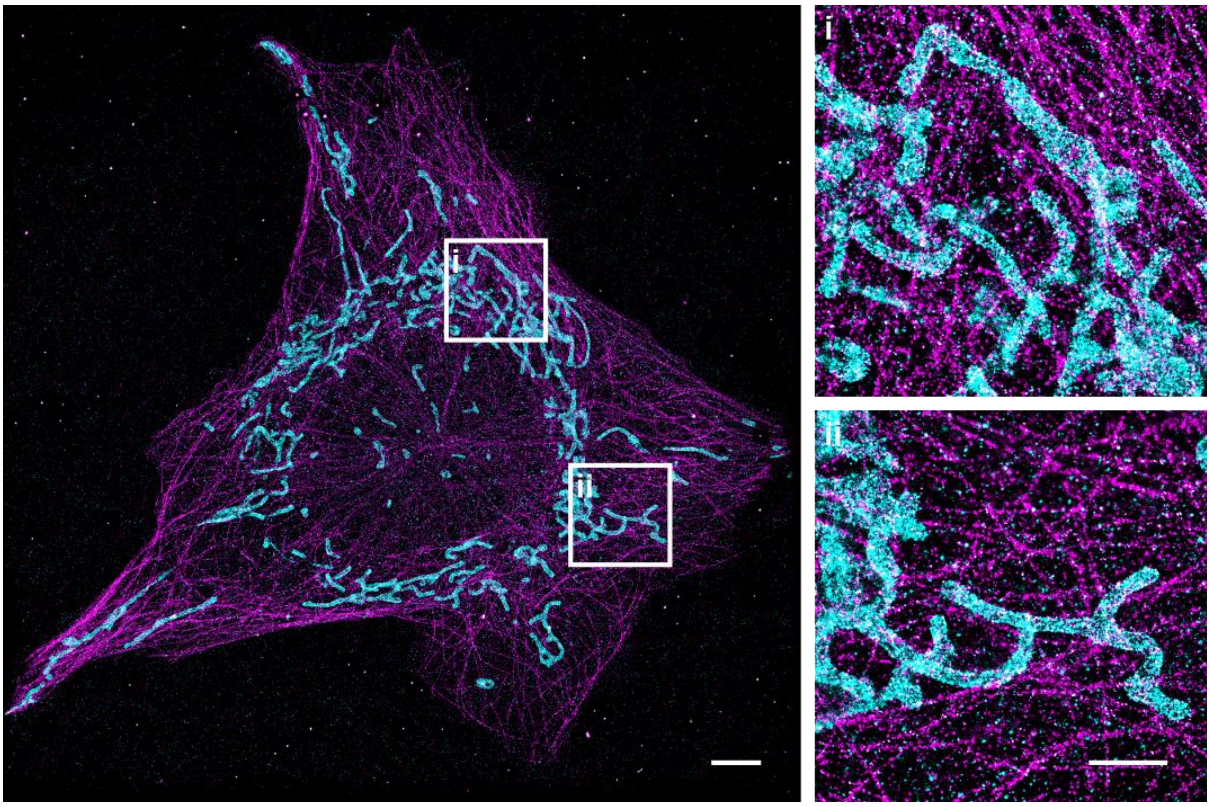


**Figure S6.** Two-target DNA-PAINT imaging of microtubules and mitochondria with caged docking strands. α-Tubulin was labeled with **cP1**$_a$ (magenta), and TOM20 with **cP1**$_b$ (cyan). 2 nM P1(9nt)-Cy3B imager strand was used for imaging. The two-target image was generated by combining images recorded before and after violet-light illumination (2 min). The NeNA (nearest-neighbor analysis) localization precision for the DNA-PAINT image of α-tubulin was 6.36 nm and for TOM20 6.77 nm. Scale bar 5 µm, zoom-ins 2 µm.

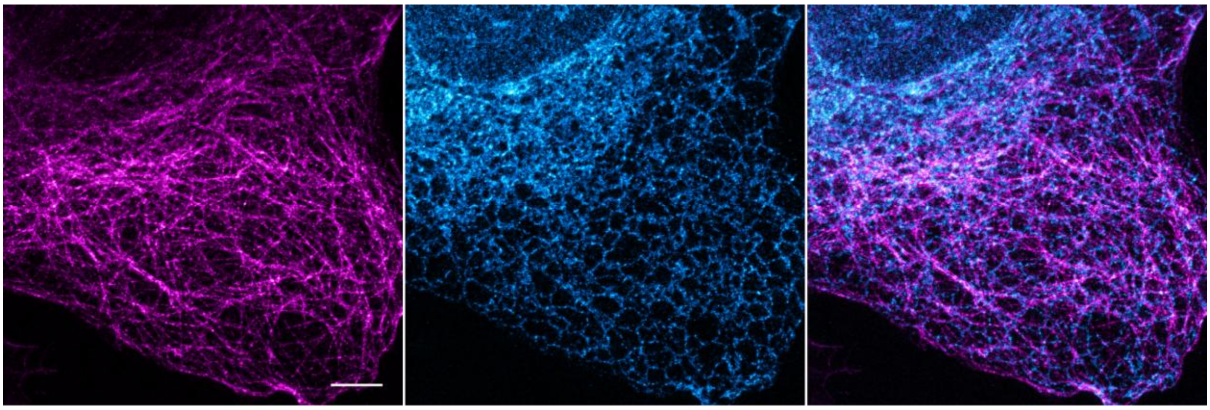

**Figure S7.** Two-target STED imaging of microtubules and ER in U-2 OS cells. α-tubulin was labeled via immunofluorescence with **cP1$_a$** docking strand (magenta) and KDEL labeled with **cP1$_b$** docking strand (cyan). Imager strand concentration was 300 nM Cy3B-P1-Cy3B. Scale bar 5 µm.

### Supporting Video

**Video S1.** Two-target confocal imaging of microtubules and mitochondria via cagePAINT. α-Tubulin was labeled with **$cP1_a$**, and TOM20 with **$cP1_b$**. At the beginning of the video, the microtubule structure is visible. Upon illumination with violet light, this structure vanishes, and the mitochondrial structure appears. The same cells as in **Figure 1D** are shown. Imager strand concentration: 100 nM P1-Cy3B. Scale bar 20 μm.

### Experimental Procedures

#### Materials and Methods

Unless otherwise stated, all reactions were performed at room temperature under an argon atmosphere. When the Schlenk technique was required, the reaction flask was heated under vacuum and evacuated, and backfilled with argon three times.

All solvents used for synthesis had a purity of ≥ 95% and were used without further purification. They were purchased in septum-sealed bottles and stored under an inert gas atmosphere with molecular sieves. Solvents for column chromatography were of technical grade, except for the phosphitylation step, for which HPLC-grade solvents were used.

Chemicals were purchased from Sigma-Aldrich (St. Louis, MO, USA), BLD Pharm (Shanghai, China), TCI Chemicals (Tokyo, Japan), Thermo Fisher (Scientific, Waltham, MA, USA), Fluorochem (Derbyshire, UK), VWR (Radnor, PA, USA), and Carl Roth (Karlsruhe, Germany).

All reactions were monitored with Thin Layer Chromatography (TLC). Therefore, the $R_f$-values of this work refer to 60 DC-Fertigfolien ALUGRAMR Xtra SIL G/UV254 TLC plates by Macherey-Nagel (Düren, Germany) with 0.2 mm silica gel coating. Detection was performed using UV light at wavelengths of 254 nm and 366 nm. If necessary, the TLCs were deactivated with a solution of 1-3% triethylamine in dichloromethane (DCM) before usage.

For common column chromatography, silica gel 60 from Macherey-Nagel and the eluent mixtures written in the experimental implementation section were used. For Flash Chromatography, a Puriflash XS420 from Interchim (Montlucon Cedex, France) and pre-packed flash columns with particle sizes of 15 and 30 µm by Interchim were used for normal-phase chromatography (PF-SiHP).

NMR spectroscopy was performed at the NMR service facility (Campus Riedberg, Goethe University Frankfurt) on the following spectrometers: Avance HD AV400 (400 MHz $^{1}$H, 101 MHz $^{13}$C), Avance III HD AV500 (500 MHz $^{1}$H, 126 MHz $^{13}$C), Avance DRX600 (600 MHz, $^{1}$H, 151 MHz $^{13}$C). Additionally, deuterated solvents were used ($CDCl_3$-$d_6$ $^{1}$H δ = 7.26, $^{13}$C δ = 77.16, DMSO-$d_6$; $^{1}$H, δ = 2.50, $^{13}$C δ = 39.52) by the company Eurisotop (Saint-Aubin, France). For the interpretation of the NMR spectra, MestReNova (Mestrelab Research, Barcelona, Spain) was utilized. The chemical shift δ is stated in ppm and the coupling constant J in Hz. The following abbreviations were used to characterize the signals' multiplicities: s (singlet), d (doublet), dd (doublet of doublets), t (triplet), q (quartet), m (multiplet).

High-resolution mass spectrometry was performed by the Mass Spectrometry Service (Campus Riedberg, Goethe University Frankfurt). The measurements were performed on a Bruker MicrOTOF qII (Bruker Corporation, Billerica, MA, USA) with sodium formate calibration segment correction.

#### Chemical Synthesis

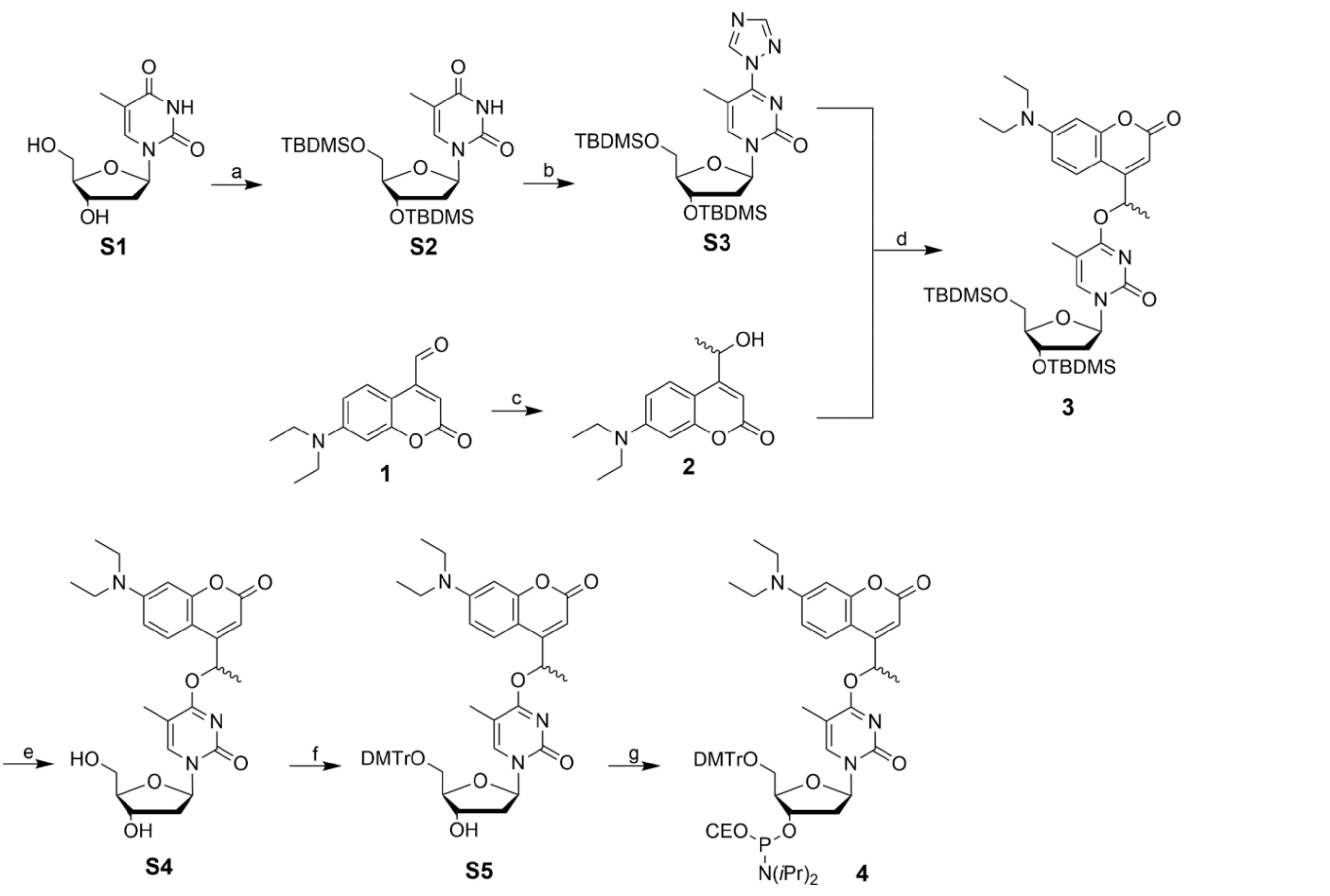

**Scheme S1.** Synthesis of Methyl-DEACM caged dT -Phosphoramidite **4.** a. dT, imidazole, TBDMS-Cl. DMF, 0 °C to rt, 20 h, quant. b. 1,2,4-triazole, $POCl_3$, triethylamine, ACN, 0 °C to rt, 19 h, quant. c. MeMgBr, THF, -78 °C to rt, 21 h, 41%. d. **S3**, DBU, ACN, rt, 20 h, 77%. e. TBAF, AcOH, THF, 0 °C to rt, 20 h, 85%. f. DMTr-Cl, DIPEA, DCM, 0 °C to rt, on, 83%. g. PN(*i*Pr)$_2$(CEO)-Cl, DIPEA, DCM, 0 °C to rt, 1.5 h, 80%.

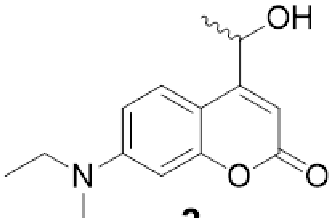

**2**

**7-(diethylamino)-4-(1-hydroxyethyl)-2H-chromen-2-one (2):** 7-(diethylamino)-2-oxo-2H-chromene-4-carbaldehyde **1** (1.00 g, 4.08 mmol, 1 eq) was dissolved in 30 mL dry THF in a Schlenk flask and cooled to -78 °C. A solution of MeMgBr (3 M in diethyl ether, 1.50 mL, 4.48 mmol, 1.1 eq) was added slowly, and the reaction mixture was stirred at -78 °C for 1 h. Afterwards, the solution was further stirred at room temperature for 1 h. The reaction was quenched by the addition of $NH_4Cl$ solution, extracted with DCM, and the solvents were removed under reduced pressure. After column chromatography (cyclohexane/ethyl acetate 1:1) **2** was afforded as a brown foam (438 mg, 41%).

$R_f$=0.4 (cyclohexane/ethyl acetate 1:1). $^1$H NMR (500 MHz, $CDCl_3$) δ = 7.42 (d, *J* = 9.0 Hz, 1H), 6.57 (d, *J* = 8.9 Hz, 1H), 6.49 (d, *J* = 2.3 Hz, 1H), 6.27 (s, 1H), 5.14 (q, *J* = 6.5 Hz, 1H), 3.40 (q, *J* = 7.1 Hz, 4H), 1.55 (dd, *J* = 6.6, 0.9 Hz, 3H), 1.19 (t, *J* = 7.1 Hz, 6H) ppm. $^{13}$C NMR (126 MHz, $CDCl_3$) δ = 163.05,

163.02, 159.80, 159.76, 156.58, 150.41, 125.15, 108.69, 106.33, 104.54, 98.05, 66.04, 44.87, 23.73, 12.58 ppm. HRMS (ESI) *m/z* calcd for $C_{15}H_{19}NO_3$ [M+H$^+$] = 262.1438 Da, found = 262.1450 Da, Δm = 0.00123, error = 4.7 ppm.

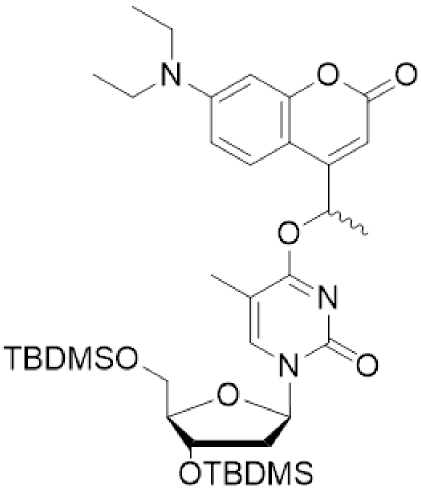


**3**

**1-((2R,4S,5R)-4-((tert-butyldimethylsilyl)oxy)-5-(((tert-butyldimethylsilyl)oxy)methyl) tetrahydrofuran-2-yl)-4-(1-(7-(diethylamino)-2-oxo-2H-chromen-4-yl)ethoxy)-5-methylpyrimidin-2(1H)-one (3):** 1-((2R,4S,5R)-4-((tert-butyldimethylsilyl)oxy)-5-(((tert-butyldimethylsilyl)oxy)methyl)tetrahydrofuran-2-yl)-5-methyl-4-(1H-1,2,4-triazol-1-yl)pyrimidin-2(1H)-one **S3** was synthesized starting from **S1** according to literature.[1] Compound **S3** (3.0 g, 5.74 mmol, 1 eq) and **2** (1.5 g, 5.74 mmol, 1 eq) were dissolved in 120 mL dry acetonitrile, and DBU (1.97 mL, 13.2 mmol, 2.3 eq) was added. The reaction was stirred under an argon atmosphere in the dark for 16 h. The solvents were removed under reduced pressure, and the precipitate was redissolved in DCM. The organic phase was washed with water and brine, followed by an additional extraction of the aqueous phase with DCM. The solvent was removed *in vacuo.* Purification by column chromatography (cyclohexane/ethyl acetate 2:1) afforded a light-yellow solid (3.15 g, 77%).

$R_f$=0.4 (cyclohexane/ethyl acetate 2:1). $^1$H NMR (400 MHz, $CDCl_3$) δ = 7.82 (d, *J* = 1.0 Hz, 1H), 7.50 (dd, *J* = 9.0, 5.4 Hz, 1H), 6.72 (d, *J* = 6.8 Hz, 1H), 6.66 – 6.59 (m, 2H), 6.30 (q, *J* = 6.4 Hz, 1H), 6.27 – 6.19 (m, 1H), 4.41 – 4.35 (m, 1H), 3.93 (ddd, *J* = 13.2, 8.5, 2.1 Hz, 2H), 3.81 – 3.74 (m, 1H), 3.42 (q, *J* = 7.1 Hz, 4H), 2.46 (tdd, *J* = 13.5, 6.2, 3.8 Hz, 1H), 2.03 (td, *J* = 4.8, 1.7 Hz, 3H), 2.01 – 1.95 (m, 1H), 1.67 (dd, *J* = 6.6, 2.2 Hz, 3H), 1.22 (t, *J* = 7.1 Hz, 8H), 0.92 (s, *J* = 2.9 Hz, 9H), 0.88 (d, *J* = 4.6 Hz, 9H), 0.11 (d, *J* = 3.3 Hz, 6H), 0.07 – 0.05 (m, 6H). ppm. $^{13}$C NMR (101 MHz, $CDCl_3$) δ = 169.02, 168.99, 156.45, 155.83, 155.77, 155.65, 140.63, 140.60, 125.55, 125.50, 104.30, 88.13, 86.69, 86.59, 71.76, 71.64, 69.24, 69.14, 62.74, 45.85, 42.47, 42.39, 26.10, 25.90, 21.30, 21.09, 18.58, 18.14, 12.57, 12.37, -4.41, -4.73, -5.21, -5.25 ppm. HRMS (ESI) *m/z* calcd for $C_{37}H_{59}N_3O_7Si_2$ [M+H$^+$] = 714.3964 Da, found = 714.3997 Da, Δm = 0.00327, error = 4.5 ppm.

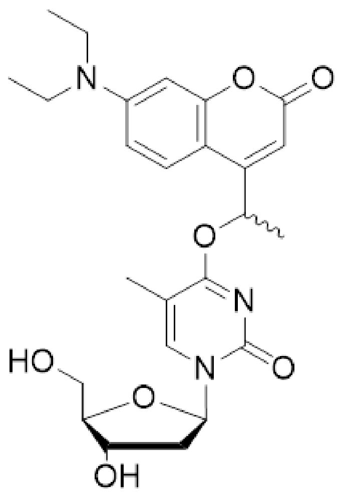


**S4**

**4-(1-(7-(diethylamino)-2-oxo-2H-chromen-4-yl)ethoxy)-1-((2R,4S,5R)-4-hydroxy-5-(hydroxymethyl)tetrahydrofuran-2-yl)-5-methylpyrimidin-2(1H)-one (S4):** Compound **3** (850 mg, 1.2 mmol, 1 eq) was taken up in 30 mL dry THF and treated with acetic acid (817 µL, 14.3 mmol, 12 eq) and a tert-butyl ammonium fluorid solution (1 M in THF, 4.8 mL, 4.76 mmol, 4 eq) at 0 °C. The reaction was allowed to warm to room temperature over 16 h. The solvents were removed under reduced pressure, followed by an extraction using DCM and $NH_4Cl$ solution. After flash chromatography (DCM to DCM/MeOH 9:1), **S4** was obtained as a yellow solid (491 mg, 85%).

$R_f$=0.51 (DCM/MeOH 95:5). $^1$H NMR (400 MHz, $CDCl_3$) δ = 7.86 (dd, *J* = 36.6, 0.9 Hz, 1H), 7.45 (dd, *J* = 9.2, 2.6 Hz, 1H), 6.62 – 6.54 (m, 2H), 6.49 (dd, *J* = 2.4, 1.4 Hz, 1H), 6.16 – 6.08 (m, 2H), 4.59 – 4.50 (m, 1H), 4.01 (dd, *J* = 6.4, 3.0 Hz, 1H), 3.94 – 3.88 (m, 1H), 3.85 – 3.78 (m, 1H), 3.40 (q, *J* = 7.1 Hz, 4H), 2.41 (t, *J* = 5.9 Hz, 2H), 2.03 (dd, *J* = 2.4, 0.8 Hz, 3H), 1.67 (d, *J* = 6.6 Hz, 3H), 1.19 (dd, *J* = 9.7, 4.4 Hz, 6H) ppm.$^{13}$C NMR (101 MHz, $CDCl_3$) δ = 169.30, 162.82, 156.64, 156.08, 155.89, 150.76, 142.22, 125.35, 108.96, 105.79, 105.03, 104.55, 97.97, 88.72, 88.18, 87.75, 71.02, 70.65, 69.73, 62.07, 44.88, 40.80, 21.26, 12.59,12.32 ppm. HRMS (ESI) *m/z* calcd for $C_{25}H_{31}N_3O_7$ [M+H$^+$] = 486.2235 Da, found = 486.2231 Da, Δm = 0.00038, error = 0.8 ppm.

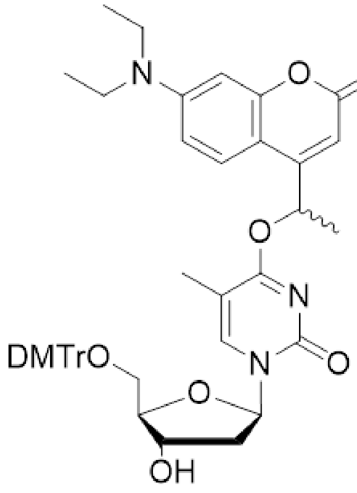


**S5**

**1-((2R,4S,5R)-5-((bis(4-methoxyphenyl)(phenyl)methoxy)methyl)-4-hydroxytetrahydrofuran-2-yl)-4-(1-(7-(diethylamino)-2-oxo-2H-chromen-4-yl)ethoxy)-5-methylpyrimidin-2(1H)-one (S5):** Compound **S4** (480 mg, 988.6 µmol, 1.0 eq) was dissolved in 12 mL dry DCM, *N,N*-Diisopropylethylamin (861 µL, 4.9 mmol, 5 eq ) was added and the mixture was cooled to 0 °C. Dimethoxytritychlorid (435 mg, 1.3 mmol, 1.3 eq) was added portion-wise, and the reaction mixture was allowed to warm to room temperature and stirred overnight in the dark under an argon atmosphere. The reaction was terminated by adding MeOH, and the solvents were removed *in vacuo*. After column chromatography (DCM/MeOH 9:1), **S5** was obtained as a yellow solid (555 mg, 83%).

$R_f$=0.6 (DCM/MeOH 92:8). $^1$H NMR (500 MHz, $CDCl_3$) δ 7.92 (dd, *J* = 4.1, 0.8 Hz, 1H), 7.45 (t, *J* = 9.3 Hz, 1H), 7.42 – 7.38 (m, 2H), 7.32 – 7.27 (m, 6H), 7.26 – 7.22 (m, 1H), 6.84 (dd, *J* = 8.8, 1.8 Hz, 4H), 6.63 – 6.56 (m, 2H), 6.51 – 6.49 (m, 1H), 6.40 – 6.33 (m, 1H), 6.16 (dd, *J* = 12.2, 0.5 Hz, 1H), 4.59 – 4.53 (m, 1H), 4.09 (d, *J* = 3.5 Hz, 1H), 3.79 (s, 3H), 3.79 (d, *J* = 0.6 Hz, 3H), 3.51 – 3.47 (m, 1H), 3.44 – 3.37 (q, *J* = 6.5 Hz, 4H), 2.60 (dddd, *J* = 20.1, 13.7, 6.1, 4.2 Hz, 1H), 2.40 (d, *J* = 13.1 Hz, 1H), 2.28 (tt, *J* = 13.0, 6.4 Hz, 1H), 1.67 (s, 1H), 1.65 (d, *J* = 6.8 Hz, 3H), 1.62 (dd, *J* = 4.6, 0.6 Hz, 3H), 1.20 (t, *J* = 7.0 Hz, 6H).ppm. $^{13}$C NMR (126 MHz, $CDCl_3$) δ = 169.16, 162.65, 162.57, 158.83, 156.65, 155.87, 155.79, 155.63, 150.69, 144.54, 140.64, 135.60, 135.52, 130.22, 128.19, 127.27, 125.34, 113.43, 108.84, 105.88, 104.85, 104.80, 98.01, 87.03, 86.49, 86.23, 71.95, 69.35, 63.32, 60.55, 55.42, 53.57, 44.87, 42.10, 29.84, 21.26, 14.34, 12.60, 11.96 ppm. HRMS (ESI) *m/z* calcd for $C_{46}H_{49}N_3O_9$ [M+H$^+$] = 788.3542 Da, found = 788.3570 Da, Δm = 0.0028, error = 3.6 ppm.

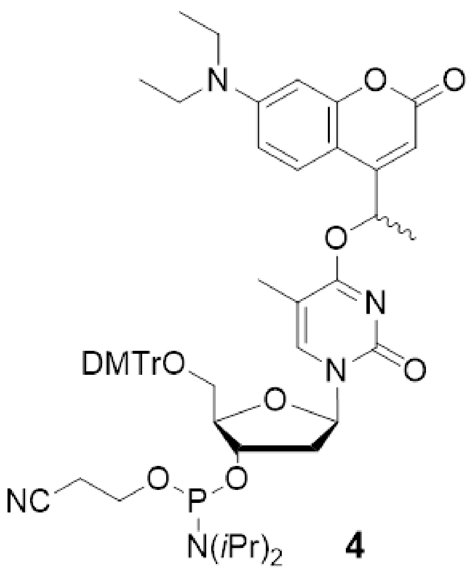


**4**

**(2R,3S,5R)-2-((bis(4-methoxyphenyl)(phenyl)methoxy)methyl)-5-(4-(1-(7-(diethylamino)-2-oxo-2H-chromen-4-yl)ethoxy)-5-methyl-2-oxopyrimidin-1(2H)-yl)tetrahydrofuran-3-yl (2-cyanoethyl) diisopropylphosphoramidite (4):** To a Schlenk flask a solution of compound **S5** (280 mg, 355.4 µmol, 1 eq) in 3.5 mL dry DCM, *N,N*-Diisopropylethylamin (310 µL, 1.8 mmol, 5 eq) and 2-Cyanethyl-*N,N*-diisopropylchlorphosphoramidit (119 µL, 533.1 µmol, 1.5 eq) were added. The reaction was stirred at room temperature for 1.5 h. The solution was diluted with DCM, and after an extraction with DCM and $NaHCO_3$, flash column chromatography (DCM to DCM/MeOH 95:5) followed. The resulting solid was co-evaporated with toluene three times, redissolved in DCM, and precipitated with *n*-hexane at -20 °C for 30 min, followed by centrifugation (20 min, 15 °C, 7830 rpm). This precipitation–centrifugation cycle was repeated three times, and compound **4** was yielded as a yellow solid (282 mg, 80%).

$R_f$=0.9 (DCM/MeOH 96:4). $^1$H NMR (500 MHz, $CDCl_3$) δ = 7.96 (ddd, *J* = 28.4, 6.2, 0.8 Hz, 1H), 7.46 (ddd, *J* = 9.6, 7.6, 2.5 Hz, 1H), 7.43 – 7.38 (m, 2H), 7.29 (qd, *J* = 8.6, 2.2 Hz, 6H), 7.26 – 7.21 (m, 1H), 6.86 – 6.81 (m, 4H), 6.64 – 6.57 (m, 2H), 6.50 (t, *J* = 2.4 Hz, 1H), 6.42 – 6.30 (m, 1H), 6.16 (dd, *J* = 8.6, 3.6 Hz, 1H), 4.69 – 4.59 (m, 1H), 4.18 – 4.14 (m, 1H), 3.79 (dd, *J* = 4.4, 1.9 Hz, 6H), 3.61 – 3.49 (m, 4H), 3.44 – 3.38 (m, 4H), 3.33 (ddd, *J* = 10.5, 8.4, 3.0 Hz, 1H), 2.75 (td, *J* = 6.1, 3.0 Hz, 1H), 2.64 – 2.58 (m, 1H), 2.41 (t, *J* = 6.3 Hz, 1H), 2.33 – 2.23 (m, 1H), 1.66 (dd, *J* = 11.0, 3.9 Hz, 6H), 1.55 (dd, *J* = 10.2, 6.7 Hz, 3H), 1.21 (t, *J* = 7.0 Hz, 6H), 1.18 – 1.13 (m, 6H), 1.03 (dd, *J* = 6.7, 4.4 Hz, 3H) ppm. $^{13}$C NMR (126 MHz, $CDCl_3$) δ = 171.29, 169.12, 162.60 , 158.83, 156.64, 155.89, 155.74, 155.70, 150.67, 144.45, 140.63, 135.48, 130.29, 128.34, 128.11, 127.28, 125.35, 117.67, 117.51, 117.02, 113.38, 108.82, 105.90, 104.93, 104.73, 98.00, 86.95, 86.46, 86.35, 85.67, 73.35, 72.52, 69.27, 63.05, 62.63, 60.53, 58.46, 58.27, 55.41, 45.44, 44.86, 43.36, 41.03 24.68, 23.07, 21.34, 21.19, 20.49, 20.27, 14.33, 12.60, 11.83 ppm. $^{31}$P NMR (202 MHz, $CDCl_3$) δ = 149.22, 149.12, 148.58, 148.51 ppm. HRMS (ESI) *m/z* calcd for $C_{55}H_{66}N_5O_{10}P$ [M+MeOH+H$^+$] = 1020.4882 Da, found = 1020.4919 Da, Δm = 0.0037, error = 3.6 ppm.

#### Oligonucleotide Synthesis

RNase-free water was used for all work involving oligonucleotides. Hence, Milli-Q water was treated with 0.1% diethyl pyrocarbonate (DEPC), stirred overnight, and autoclaved before usage.

Oligonucleotide synthesis was performed on an ABI 392 DNA/RNA synthesizer (Applied Biosystems, Thermo Fisher Scientific, Waltham, MA, USA) at a 1 µmol scale. 0.3 M 5-benzylmercaptotetrazole (BTT) in anhydrous acetonitrile (emp Biotech, NJ, USA) was used as the activator, phenoxyacetic anhydride and pyridine in tetrahydrofuran 5:10:85 (Capping A) (emp Biotech, NJ, USA), and 10%-1-methylimidazole in tetrahydrofuran (Capping B) (emp Biotech, NJ, USA) were used as UltraMild capping agents. Oxidizing (ABI) (J.T. Baker, Avantor, Radnor, PA, USA) was used as the oxidizing reagent, and 3% trichloroacetic acid in dichloromethane was used as a deblocking reagent (emp Biotech, NJ, USA).

Commercially available DNA amidites were all 5'-DMTr-protected and purchased from LGC Biosearch Technologies, Hoddesdon, UK (dC (Ac) CE-phosphoramidite, dG (iPr-Pac) CE-phosphoramidite, dT CE-phosphoramidite, 5'-MMTr C6 Amino Modifier phosphoramidite) and Sigma-Aldrich (dA (tac) CE-phosphoramidite). For solid support, dG (iPr-Pac) 1000 Å from LGC Biosearch Technologies was used.

All syntheses were performed in an MMTr-On mode with coupling times for all commercially available DNA phosphoramidites of 30 s and for the in-house synthesized phosphoramidite **4** of 15 min.

The oligonucleotides listed in **Table S1** were synthesized using the in-house synthesized phosphoramidite **4** and MMTr-C6 amino modifier (**Mod**, Chem Genes, Wilmington, MA, USA) to post-synthetically introduce an azide label (chemical structures are shown in **Figure S8**).

**Table S1.** Overview of the oligonucleotide sequences synthesized in this work. Mod. = 5'-MMTr C6 amino modifier.

| | **sequence** |
|---|---|
| **cP1$_a$** | 5'- Mod. TTA TAC ATC TAT TT**4** AGA **4**GT A**4**A AG -3' |
| **cP1$_b$** | 5'- Mod. TTA **4**AC A**4**C **4**AG -3' |

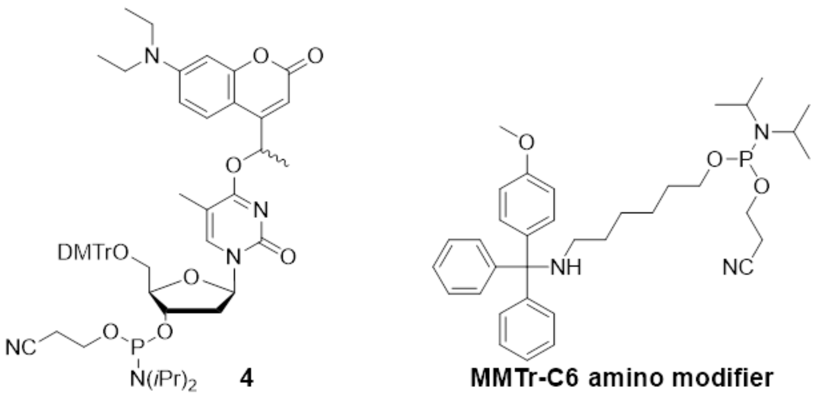

**Figure S8:** Chemical structures of in-house synthesized phosphoramidite **4** and MMTr amino modifier used in this work.

After synthesis, the columns were dried *in vacuo* for a few hours before the cyanoethyl groups were removed using 20% diethylamine in acetonitrile (emp Biotech, NJ, USA) for 15 min before drying them again. This was followed by cleavage of the solid support with aqueous ammonia (32%, Sigma-Aldrich, St. Louis, MO, USA) at 25 °C, 800 rpm for 3 hours. After spinfiltration, the solvent was removed at 4 °C in a vacuum concentrator (SpeedVac$^{TM}$, Thermo Fisher Scientific, Waltham, MA, USA),

#### Purification

The MMTr-on oligonucleotides were purified by RP-HPLC with an Agilent 1200 (Agilent Technologies, Santa Clara, CA, USA) using a Waters XBridge Peptide BEH C18 OBD Prep Column, 300 Å, 5 µm, 10 x 250 mm, 3.5 mL/min, 60 °C. As the buffer system, 400 mM hexafluoroisopropanol (Fluorochem, Derbyshire, UK), 16.3 mM trimethylamine (Sigma-Aldrich, St. Louis, MO, USA), pH 7.9, and methanol were used. A gradient from 5 to 100% methanol in 45 min was used. After separation, the solvent was removed *in vacuo* at 4 °C.

#### Azide Labeling

31.73 nmol of each oligo without MMTr was dissolved in 79.3 µL borate buffer (0.1 M sodium tetraborate (Sigma-Aldrich, St. Louis, MO, USA), pH 8.4) and treated with 67.6 eq (12.69 µL) azido butyric NHS ester (Lumiprobe, Hannover, Germany) dissolved in dry DMF. The solution was then incubated at 25 °C, 800 rpm for 18 h. The solvents were removed at 4 °C under reduced pressure and further purified with an Agilent 1200 using a Waters XBridge Peptide BEH C18 OBD Prep Column, 300 Å, 5 µm, 10 x 250 mm, 3.0 mL/min, 60 °C. As the buffer system, 400 mM hexafluoroisopropanol, 16.3 mM triethylamine, pH 7.9, and methanol were used. A gradient from 5 to 100% methanol in 45 min was used. After separation, the solvent was removed *in vacuo* at 4 °C.

#### Sample Preparation for Antibody Labeling

Remaining RP-HPLC buffer ions were removed by using a 1-kDa cut-off membrane filter (Microsep Advance Centrifugal Devices with Omega Membrane 1 kDa, Pall Corporation, Port Washington, NY, USA). Before usage, each filter was washed five times with 4.5 mL DEPC-water and three times with 0.3 M NaCl at 7197 rcf, 15 °C for 20 min. The filter was loaded with the oligonucleotide dissolved in 400 µl DEPC-water, and 4 ml 0.3 M NaCl was added. The mixture was centrifuged at 7197 rcf, 15 °C for 20 min. This process was repeated three times. The remaining sodium ions were removed using the same filters using DEPC-water as an eluent. This process was performed three times before removing the solvent at 4 °C in a vacuum centrifuge. Finally, the sample was lyophilized overnight.

#### Characterization

Analytical RP-HPLC was performed on an Agilent 1200 equipped with a Waters XBridge Peptide BEH C18 OBD column, 300 Å, 3.5 µm, 4.6 x 250 mm, 0.7 mL/min, 60 °C. As the buffer system, 400 mM hexafluoroisopropanol, 16.3 mM triethylamine, pH 7.9, and methanol were used. A gradient from 0 to 100% methanol in 30 min was used. Purity and identity of the oligos were confirmed by LC-MS (Orbitrap Exploris 120, Thermo Fisher Scientific, Waltham, MA, USA) equipped with an ACQUITY$^{TM}$ Premier Peptide BEH C18 column, 300 Å, 1.7 µm, 2.1 x 150 mm, 0.2 mL/min, 60 °C with a buffer system consisting of 400 mM hexafluoroisopropanol, 16.3 mM triethylamine, pH 7.9, and methanol. The gradient was from 5 to 100% methanol in 20 min. The observation window was from 35 to 85% methanol in 7 min.

The analytical HPLC spectra of the purified oligonucleotide strands **cP1$_a$** and **cP1$_b$** are shown in **Figures S9** and **S10**.

#### Analytical HPLC Spectra

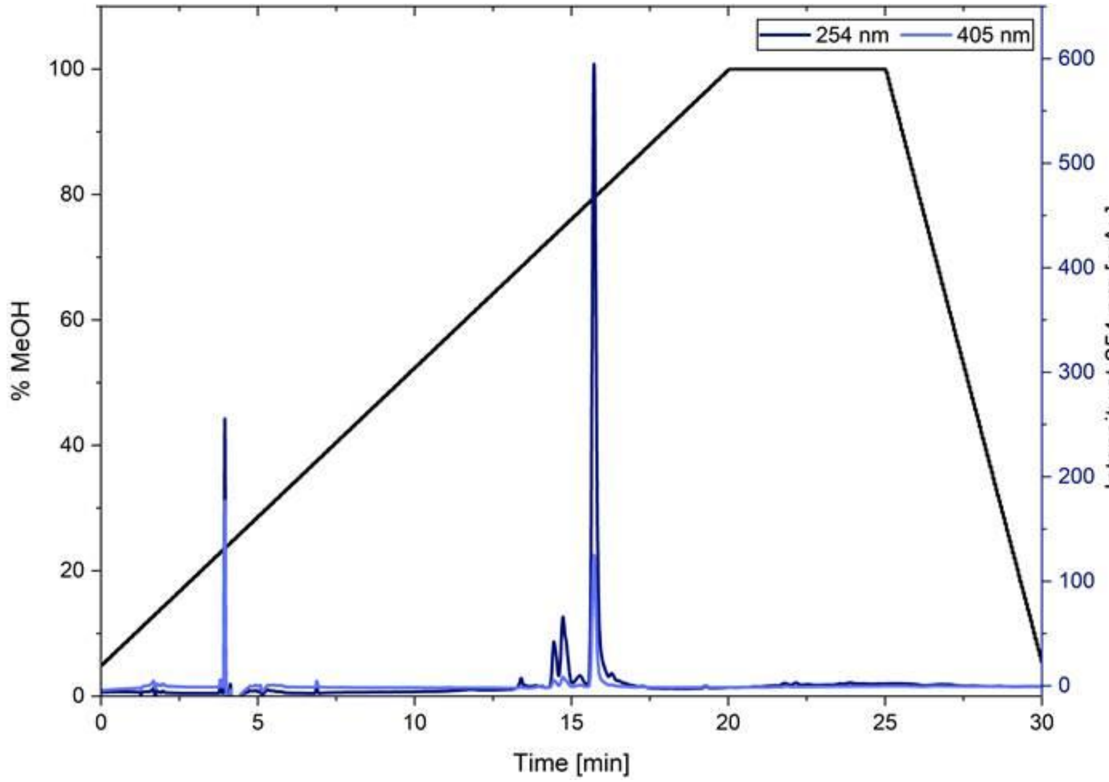


**Figure S9.** Analytical HPLC chromatogram of **cP1$_a$**. The 254 nm and 405 nm traces are shown. The peak at 3.9 min represents an injection peak, and at 15.7 min **cP1$_a$** is shown.

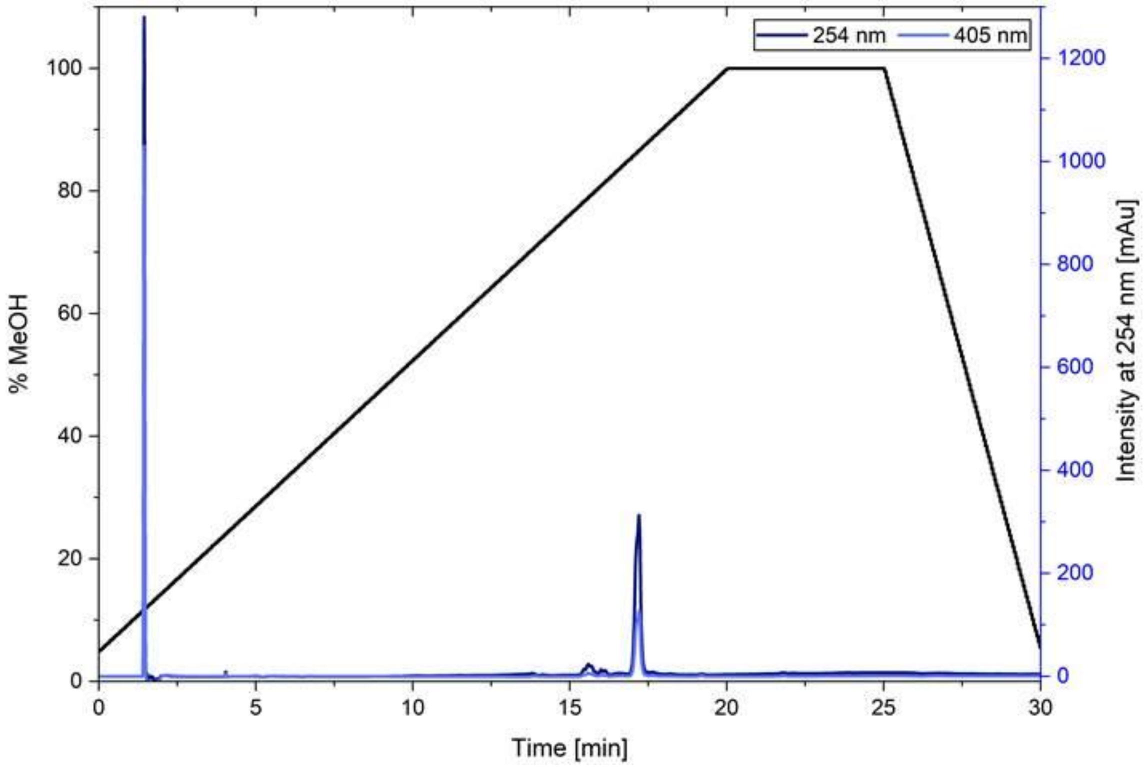


**Figure S10.** Analytical HPLC chromatogram of **cP1**$_b$. The 254 nm and 405 nm traces are shown. The peak at 1.4 min represents an injection peak, and at 17.2 min **cP1**$_b$ is shown.

Mass spectrometry confirmed the masses of both oligonucleotides as listed in **Table S2**. The corresponding mass spectra are shown in **Figures S27** and **S28**.

**Table S2.** Calculated and found masses of **cP1**$_a$ and **cP1**$_b$.

| | **calculated mass (exact) [Da]** | **found mass [Da]** |
|---|---|---|
| **cP1**$_a$ | 8990.8738 | 8988.3369 |
| **cP1**$_b$ | 4639.1581 | 4636.8755 |

#### Photochemical Characterization

The photolysis experiments of **cP1**$_a$ and **cP1**$_b$ were performed using a 405 nm LED and a driver at I = 1000 mA (both Thorlabs, Bergkirchen, Germany); this was measured to be equivalent to P = 19.8 mW. Of each oligo, 2 nmol (c = 2.5 µM) were diluted in 1× PBS buffer (pH 7.4), and uridine (c = 100 µM) was used as an internal standard. The solution was irradiated for 0 s to 600 s, and RP-HPLC was performed on an Agilent 1200 equipped with a Waters XBridge Peptide BEH C18 OBD column, 300 Å, 3.5 µm, 4.6 x 250 mm, 0.7 mL/min, 60 °C. As the buffer system, 400 mM hexafluoroisopropanol, 16.3 mM triethylamine (B), pH 7.9, and methanol (A) were used. A non-linear gradient from 0 to 100% methanol in 28 min was used (0-2 min 5% A, 2-4 min 5-20% A, 4-24 min 20-60% A, 24-28 min 60-100% A, 28-34 min 100% A, 34-38 min 100-5% A, 38-46 min 5% A).

The RP-HPLC chromatograms of the photolysis of oligonucleotide strands **cP1**$_a$ and **cP1**$_b$ are shown in **Figures S1** and **S2.**

#### Labeling of Antibodies with Photocaged Docking Strands

Antibody labeling with photocaged docking strands was accomplished as previously published.[2] All steps involving photocaged DNA were carried out with minimal light exposure, using complete darkness whenever possible. Antibodies were kept on ice throughout the procedure. Goat anti-mouse (Jackson

ImmunoResearch, West Grove, PA, USA, #115-005-003, lot 163693) and goat anti-rabbit antibodies (Jackson ImmunoResearch, #111-005-003, lot 161699) were concentrated and buffer-exchanged into 1× phosphate-buffered saline (PBS) (#14190094, Gibco, Thermo Fisher Scientific, Waltham, MA, USA) using Amicon Ultra centrifugal filters (MWCO 100 kDa; Millipore, Sigma-Aldrich, St. Louis, MO, USA). Following washes and centrifugation steps, antibody concentrations were adjusted to >1.5 mg/mL.

For crosslinking, antibodies were reacted with DBCO-sulfo-NHS ester (#762040, Sigma-Aldrich) at a 10-fold molar excess and incubated 90 min at 4°C, slowly rotating, protected from light. Excess linker was removed using Zeba spin desalting columns (#89882, Thermo Fisher Scientific), and the conjugates were concentrated to >1.5 mg/mL. DNA–antibody conjugation was performed by mixing azide-modified DNA docking strands with crosslinker-antibody conjugates at a 5:1 molar ratio and incubating overnight at 4°C, slowly rotating. Excess DNA was removed via centrifugal filtration (Amicon Ultra, MWCO 100 kDa) with PBS washes. Conjugates were collected, adjusted to about 100 µL per sample, quantified by Nanophotometer (Implen GmbH, München, Germany), and stored at 4°C in the dark.

#### Immunofluorescence

The human osteosarcoma cell line U-2 OS (#300364, CLS Cell Lines Service GmbH, Eppelheim, Germany) was cultured in high-glucose DMEM/nutrient mixture F-12 (DMEM/F12) (#21041033, Gibco, Thermo Fisher Scientific) with 1% GlutaMAX (#35050-038, Gibco), penicillin (100 unit/mL) and streptomycin (100 µg/mL; #15140122, Gibco), and 10% fetal bovine serum (FBS) (#35-079-CV, Corning Inc., Corning, NY, USA) at 37 °C and 5% $CO_2$ in an automatic $CO_2$ incubator (Model C 150, Binder GmbH, Tuttlingen, Germany). Cells were seeded at $2 \times 10^4$ cells per well in 8-well chamber slides (#94.6170.802, Sarstedt, Nümbrecht, Germany) coated with 15 µg/mL fibronectin (#F08995, Merck, Sigma-Aldrich).

Cells were fixed with 3% methanol-free formaldehyde (FA) (#28908, Thermo Scientific) and 0.1% glutaraldehyde (GA) (#G5882, Sigma-Aldrich) in 1× PBS for 15 min at room temperature. After one wash with 1× PBS, residual aldehydes were quenched with ~0.1% sodium borohydride ($NaBH_4$) (#452882, Sigma-Aldrich) for 7 min, followed by three PBS washes. Cells were blocked and permeabilized with 0.25% Triton X-100 (#T8787, Sigma-Aldrich) and 3% IgG-free bovine serum albumin (BSA) (3737.2, Roth, Karlsruhe, Germany) in 1× PBS (blocking buffer) for 10 min at room temperature. Primary antibodies (**Table S3**) were diluted in blocking buffer and applied to cells for 2 h at room temperature with gentle shaking. After three washes with PBS, self-labeled secondary antibodies (1:100 in blocking buffer) were applied for 1 h at room temperature with gentle shaking, followed by three PBS washes. Cells were post-fixed with 4% methanol-free FA in 1× PBS for 10 min at room temperature and washed three times with PBS. Gold nanoparticles (90 nm, Nanopartz, Loveland, CO, USA) were used as fiducial markers for single-molecule localization microscopy. First, gold nanoparticles were sonicated for 10 min, diluted 1:4 in 1× PBS, sonicated for an additional 10 min, and added to cells for 15 min. Wells were rinsed twice with PBS. Labeled cells were stored at 4°C in 1× PBS containing 0.05% $NaN_3$ (#S2002, Sigma-Aldrich), wrapped in Parafilm and aluminum foil to protect from light.

**Table S3.** Primary antibodies used in this study. For each antibody, the dilution used for immunostaining, the catalog number, and the company are given.

| antibody | dilution | cat. no. | company |
|---|---|---|---|
| mouse anti-α-tubulin | 1:50 | #32-2500 | Invitrogen (Thermo Fisher Scientific) |
| rabbit anti-TOM20 | 1:500 | #11802-1-AP | Proteintech (Rosemont, IL, USA) |
| rabbit anti-vimentin | 1:500 | #ab92547 | abcam (Cambridge, UK) |
| rabbit anti-KDEL | 1:200 | #ab176333 | abcam |
| mouse anti-vimentin | 1:500 | #ab8069 | abcam |
| rabbit anti-lamin B1 IgG | 1:100 | #ab16048 | abcam |

#### Confocal Laser Scanning Microscopy

Confocal imaging was performed using a Leica DMi8 inverted microscope equipped with a Leica TCS SP8 scanhead and a HC PL APO CS2 63×/1.40 oil immersion objective (Leica Microsystems, Wetzlar, Germany). Data were recorded using the Leica Application Suite X Software (v3.5.9.26787). For imaging, P1-Cy3B imager strand (9 nt: TAGATGTAT-Cy3B) (metabion, Planegg, Germany), P1-ATTO 643 imager strand (9 nt: TAGATGTAT-AT643) (biomers, Ulm, Germany), or P1-ATTO 655 imager strand (11 nt: TAGATGTATAA-AT655) (Eurofins Genomics Germany, Ebersberg, Germany) was used. For all dyes, an imaging buffer consisting of 0.5 M NaCl and 1 mM EDTA in 1× PBS was used. For ATTO 643, imaging, 1 mM Trolox was additionally included as a redox system. For Cy3B imaging, an oxygen scavenging system composed of 10 nM protocatechuate-3,4-dioxygenase (PCD) and 2.5 mM protocatechuic acid (PCA), as well as Trolox (1 mM), were included (all from Sigma-Aldrich). It has previously been shown that the use of an oxygen scavenging system in combination with a redox system reduces damage induced by reactive oxygen species, thereby preventing premature photobleaching of fluorophores and depletion of docking strands.[3,4]

The following imaging parameters were used for all images: 16-bit depth, a pixel size of 90 nm, a scan speed of 100 Hz, and a pixel dwell time of 7.69 µs. The pinhole diameter was adjusted individually for each emission wavelength (ATTO 488: 87.3 µm, Cy3B/MitoTracker Orange: 95.5 µm, ATTO 643: 53.6 µm, ATTO 655: 107.1 µm). For fluorophore excitation, the 496 nm line of the Argon ion laser (ATTO 488: 1%), the 561 nm diode laser (Cy3B: 1%, MitoTracker Orange: 0.1%), and the 633 nm HeNe laser (ATTO 643: 5%, ATTO 655: 10%) were used. For uncaging, 405-nm light at the lowest possible intensity (0.01%, corresponding to 2.72 µW before the objective and an irradiance around 8.7 kW/cm² in the sample) was used, and 5 to 10 frames were recorded (radiant exposure: ~0.3-0.6 J/cm²). Additional details on the acquisition parameters are provided in **Table S4**. Composite images of **cP1$_a$** and **cP1$_b$** were generated by combining image data acquired before and after violet-light illumination. For spatially confined uncaging, a region of interest was selected, and 405-nm light was applied exclusively to this region.

**Table S4.** Acquisition parameters for confocal measurements using the Leica SP8 microscope.

| Fig. | $\lambda_{ex}$ / nm | $\lambda_{em}$ / nm | detector (gain) | size / px | optical zoom | line average | line accu | pinhole / µm |
|---|---|---|---|---|---|---|---|---|
| 1 | 561 (1%) | 580-700 | HyD (50) | 1024x1024 | 2 | 2 | - | 95.5 |
| 2A | 496 (1%)<br>561 (0.1%)<br>633 (10%) | 510-551<br>571-650<br>660-778 | PMT (700 V)<br>HyD (50)<br>PMT (700 V) | 1024x1024 | 2 | - | 2 | 87.3<br>95.5<br>107.1 |
| S3 | 561 (1%) | 580-700 | HyD (50) | 1024x1024 | 2 | 2 | - | 95.5 |
| S4 | 561 (1%) | 580-700 | HyD (50) | 1024x1024 | 2 | 2 | - | 95.5 |
| S5 | 633 (5%) | 660-770 | HyD (20) | 2048x2048 | 1 | - | 4 | 53.6 |

#### DNA-PAINT Imaging

DNA-PAINT imaging was performed on an N-STORM microscope (Nikon, Düsseldorf, Germany) using a 561 nm laser (0.24 kW/cm$^2$) for excitation of Cy3B in HILO (highly inclined and laminated optical sheet) illumination mode. The P1-Cy3B imager strand (9 nt) was used at a concentration of 2 nM in imaging buffer containing the oxygen scavenging and redox systems described above. For data acquisition, the following parameters were set in µManager[5] (v1.4.20): image size 512x512 px, exposure time 150 ms, 30,000 frames (channel 1)/20,000 frames (channel 2), EM gain 100 (**Figure 2**)/150 (**Figure S6**), preamp gain 3, readout rate 17,000 MHz, activated frame transfer. After the first target labeled with cP1$_a$ docking strand, photocages were removed by illuminating the cell for 2 min with 405-nm light with an irradiance of 0.005 kW/cm² (radiant exposure: ~600 J/cm²). Subsequently, the second target was imaged with the same imager strand.

DNA-PAINT movies were processed with the Picasso software (v0.8.8).[2] Single-molecule localization was performed in the Picasso *Localize* module using the following settings: box side length 7, min. net gradient 70,000, EM gain 100/150, baseline 79.7, sensitivity 4.78, quantum efficiency 0.98, pixel size 157 nm. Fitting was carried out using maximum-likelihood estimation (integrated Gaussian) with a convergence criterion of 0.001 and a maximum of 1,000 iterations. After drift correction in the *Render* module using RCC (segmentation: 1000), the localization data were filtered by photon count (0-50,000), sx/sy (0.5-2 px), and lpx/lpy (0-0.2 px). The experimental localization precision was determined using a nearest-neighbor analysis[6] implemented in Picasso *Render*.

#### STED Microscopy

STED imaging was performed on a Leica Stellaris STED microscope (Leica Microsystems, Germany) using a HC PL APO CS2 100x/1.40 oil immersion objective.

For STED image acquisition, samples were excited using a pulsed white light laser with a 562 nm wavelength (8%) and depleted using a 775 nm pulsed laser (20%; ~90 MW/cm²), which had a 2D donut point spread function and an excitation delay of 300 ps. Fluorescence was collected in the spectral range of 582-744 nm using a hybrid detector (HyD). The images were acquired with a pinhole of 1.0 AU, line accumulation of 2, pixel dwell time of 7.6 µs, and an isotropic pixel size of 37.8 nm.

For uncaging, the sample was irradiated within the image boundaries using a white light laser with a 405 nm wavelength (1%). For this, a frame accumulation of 4, a line accumulation of 2, a pixel dwell time of 7.6 µs, and an isotropic pixel size of 37.8 nm were used.

Before STED imaging, a buffer was added containing Cy3B-P1-Cy3B (9 nt: Cy3B-TAGATGTAT-Cy3B) (Eurofins Genomics Germany, Ebersberg, Germany) imager strands (300 nM), PCA/PCD, and Trolox in 1× PBS. For two-target STED imaging, an initial STED image was acquired, followed by uncaging, and then a second STED image was recorded.

### Appendix

#### NMR Spectra

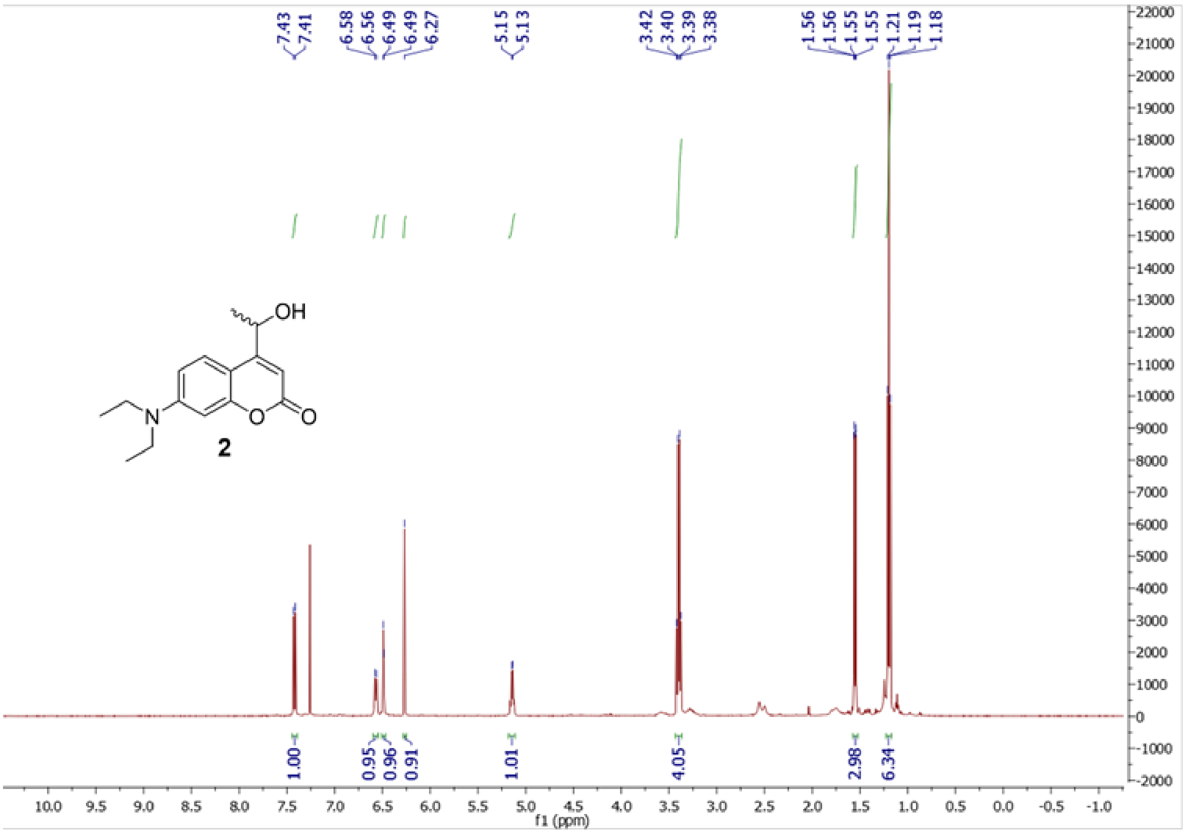


**Figure S11.** $^{1}$H NMR of **2**.

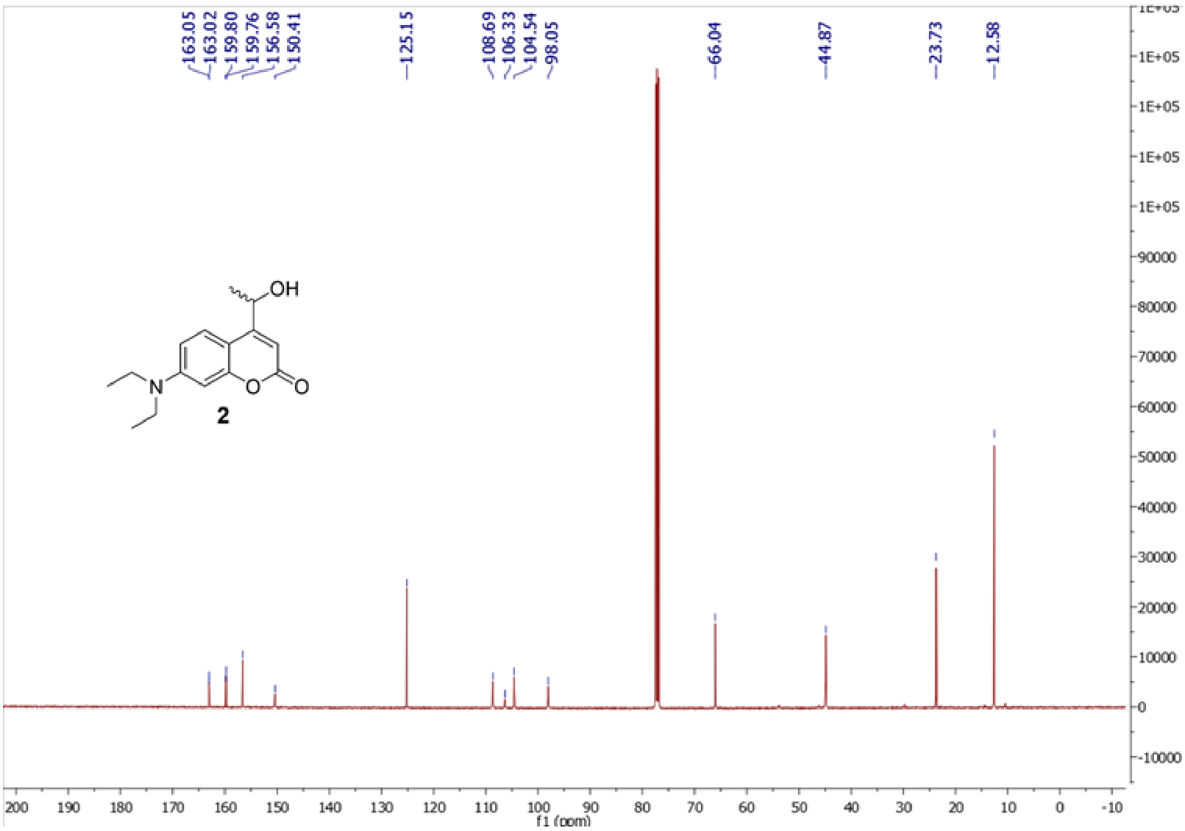


**Figure S12.** $^{13}$C NMR of **2**.

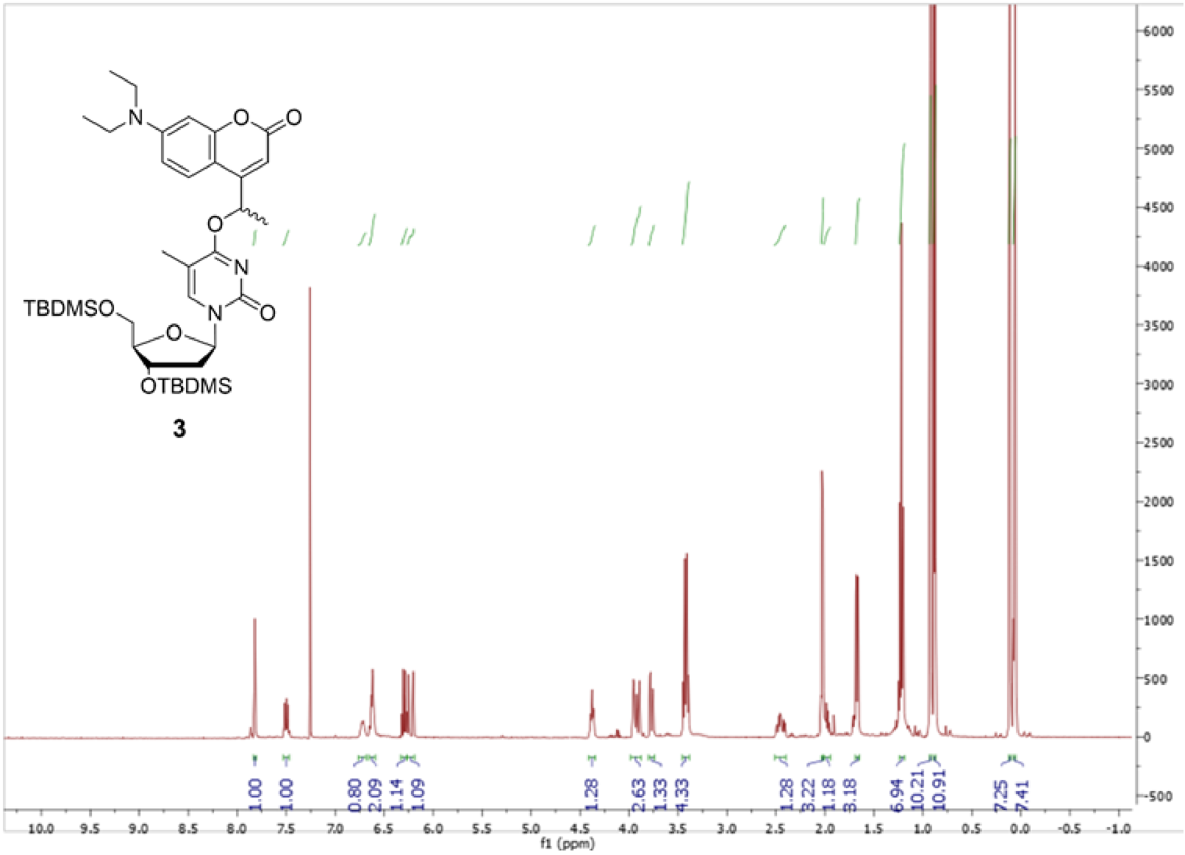

**Figure S13.** $^{1}H$ NMR of **3**.

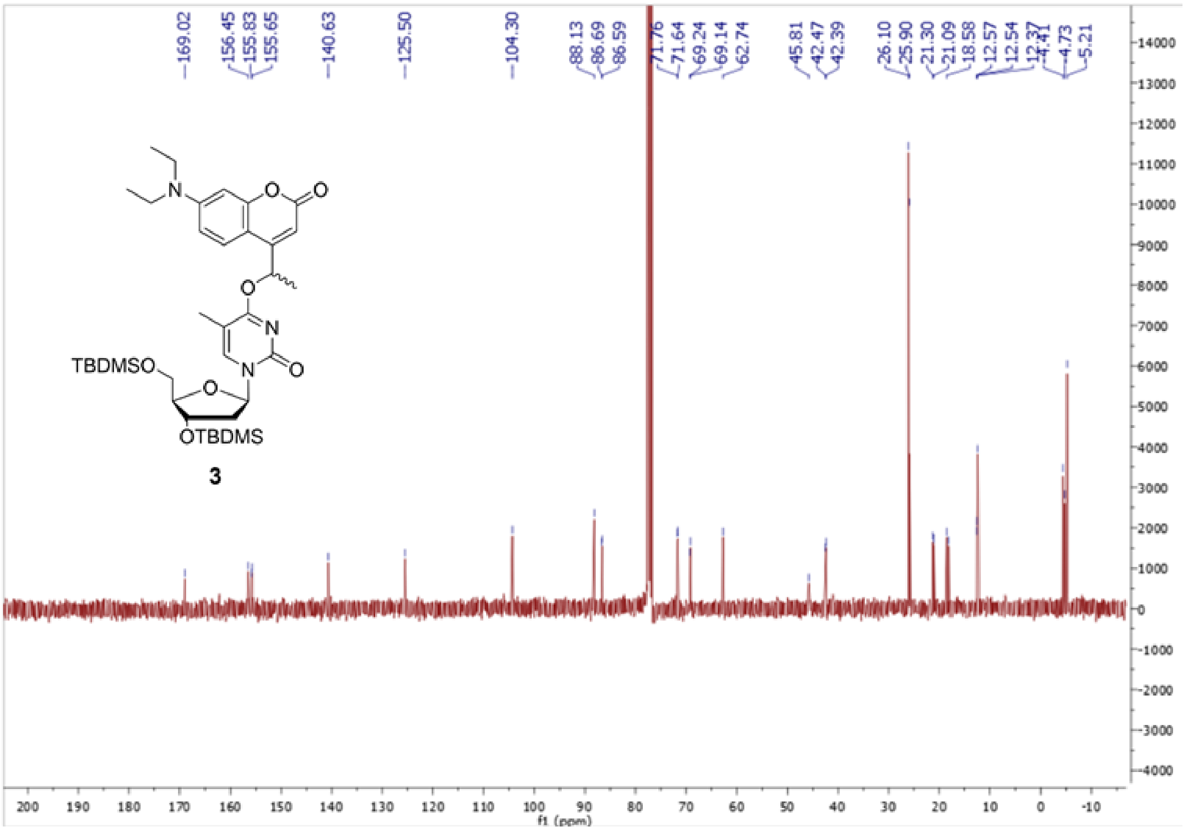

**Figure S14.** $^{13}C$ NMR of **3**.

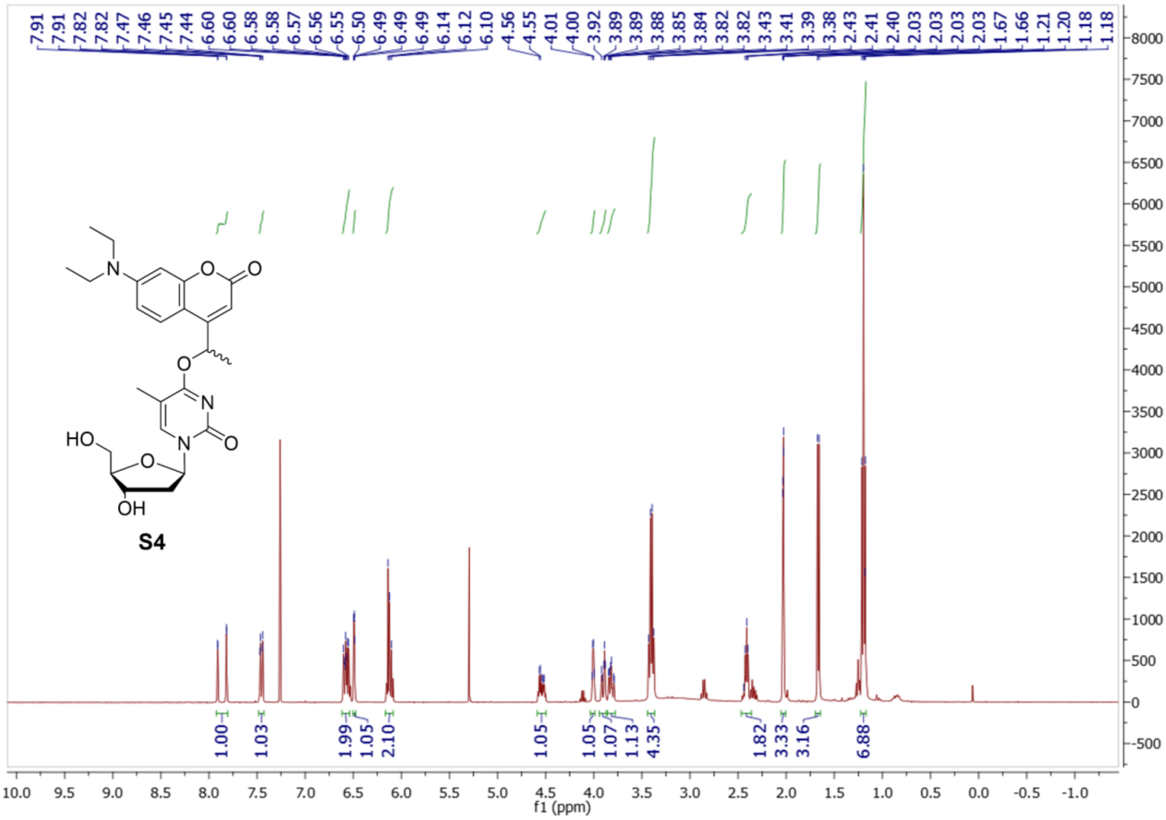

**Figure S15.** $^{1}$H NMR of **S4**.

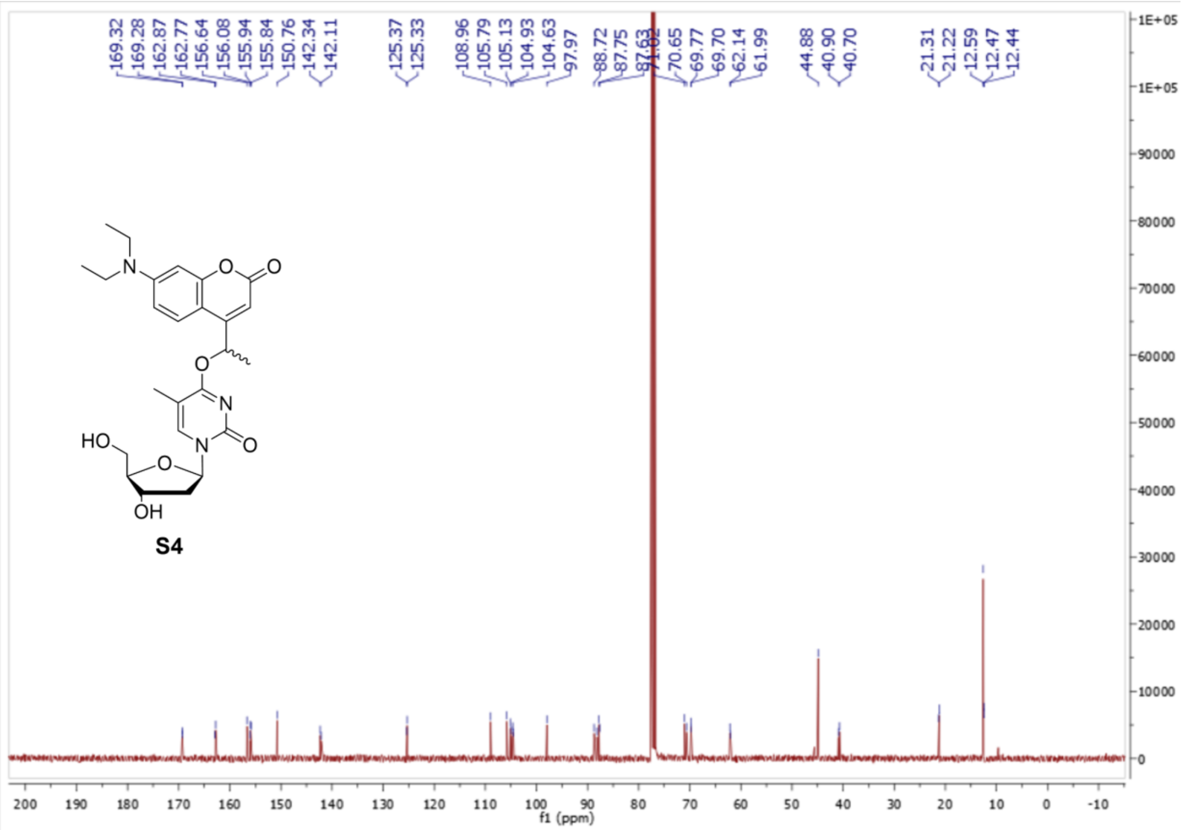

**Figure S16.** $^{13}$C NMR of **S4.**

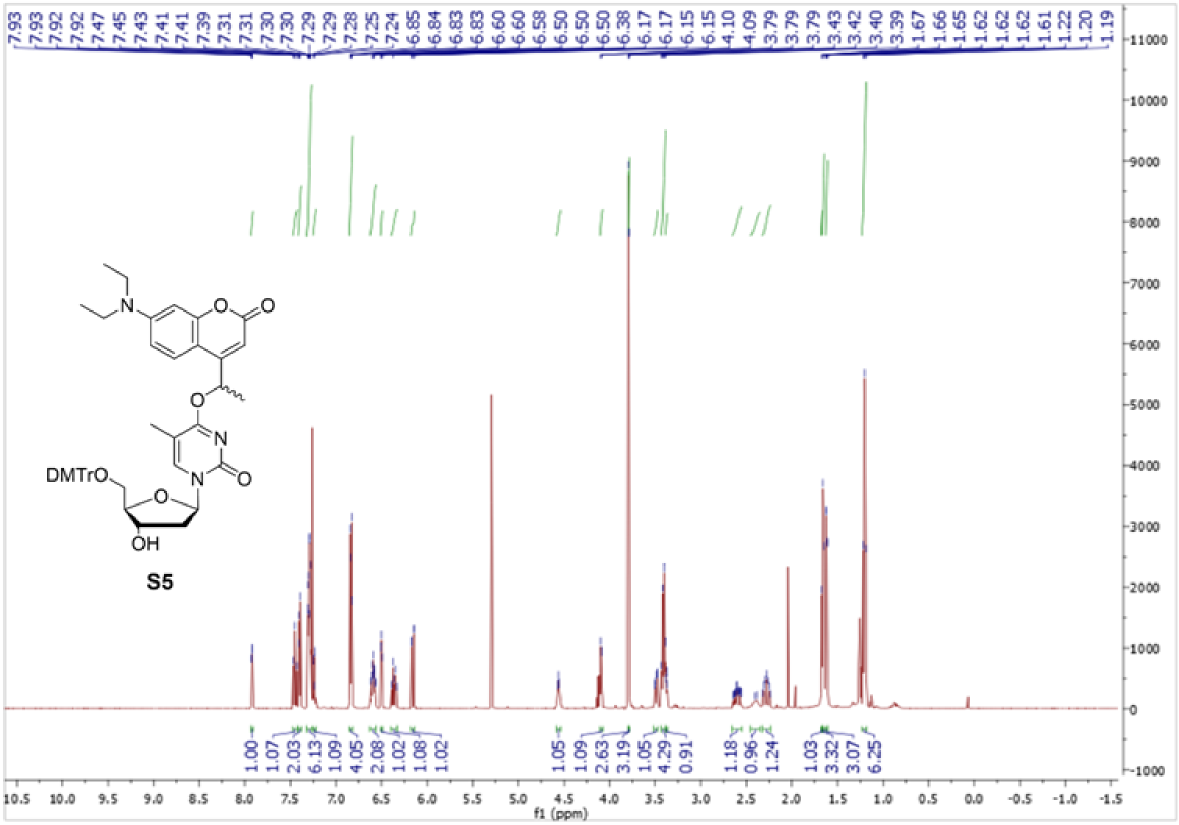

**Figure S17.** $^{1}H$ NMR of **S5**.

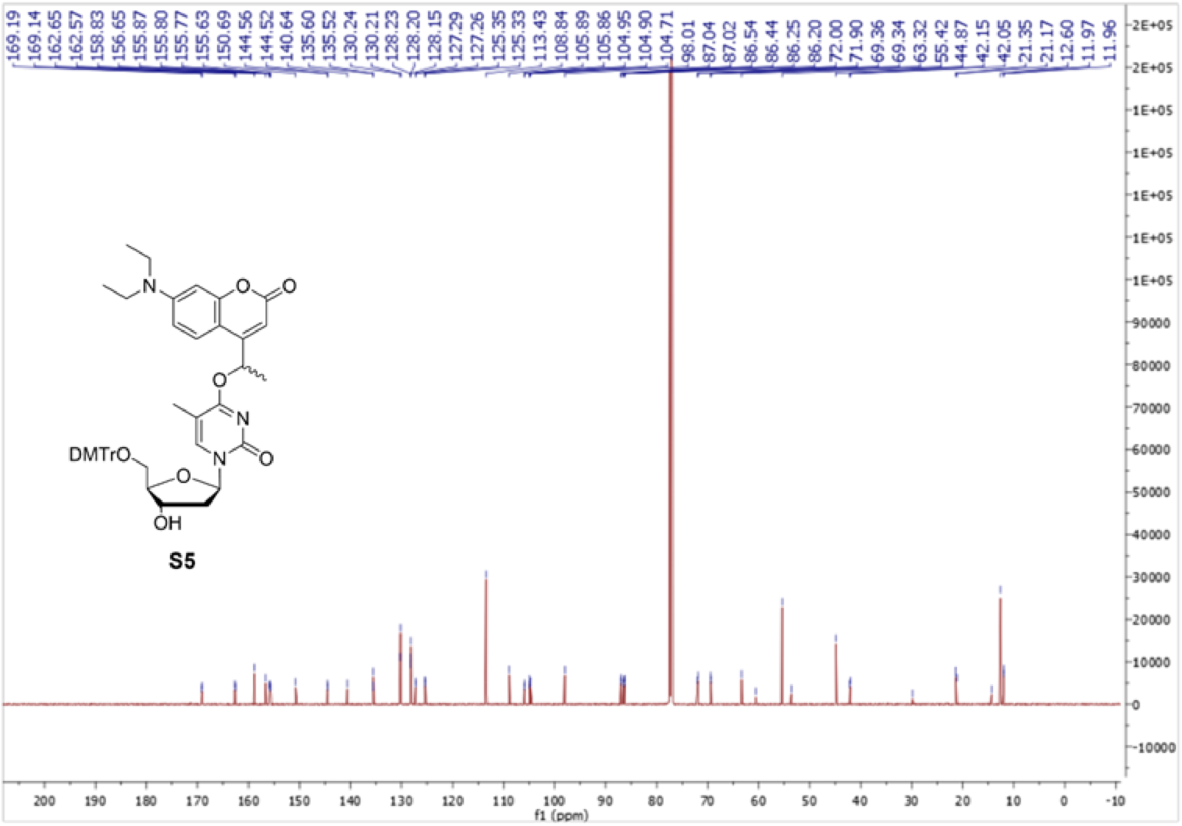

**Figure S18.** $^{13}C$ NMR of **S5**.

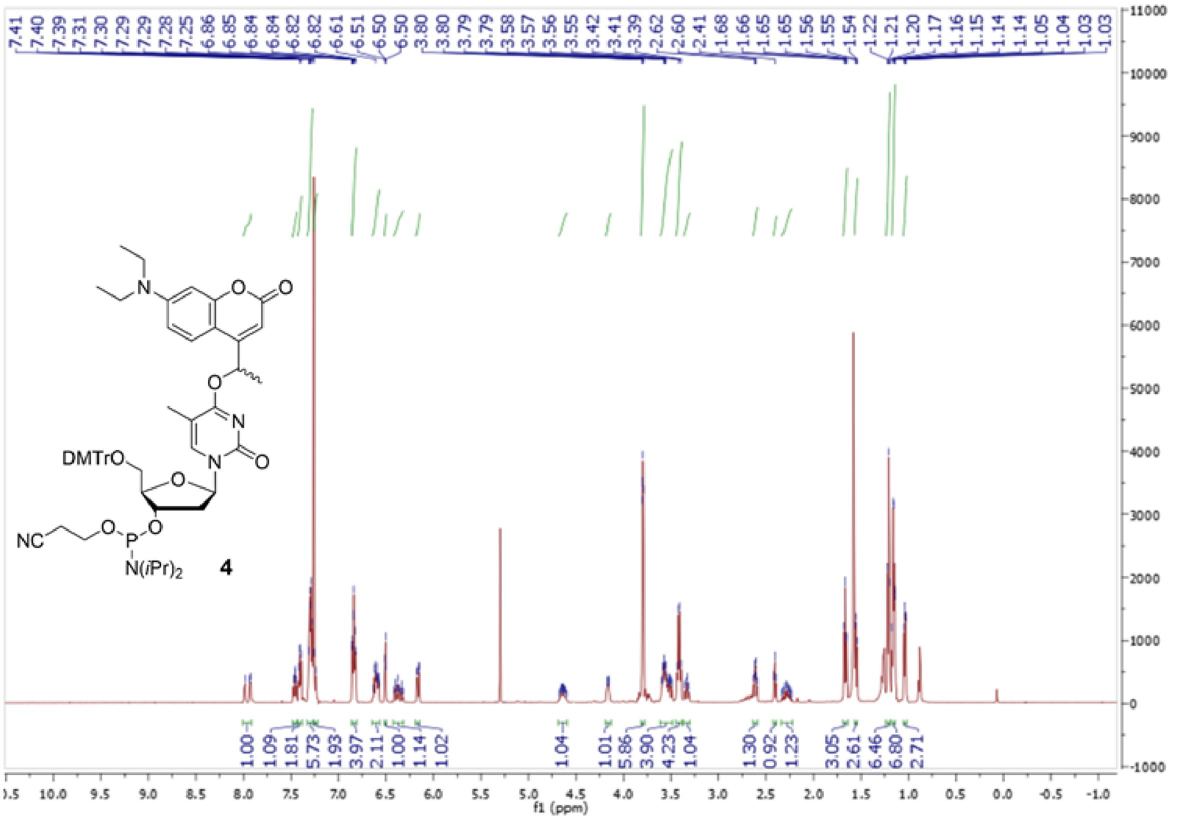

**Figure S19.** $^{1}H$ NMR of **4**.

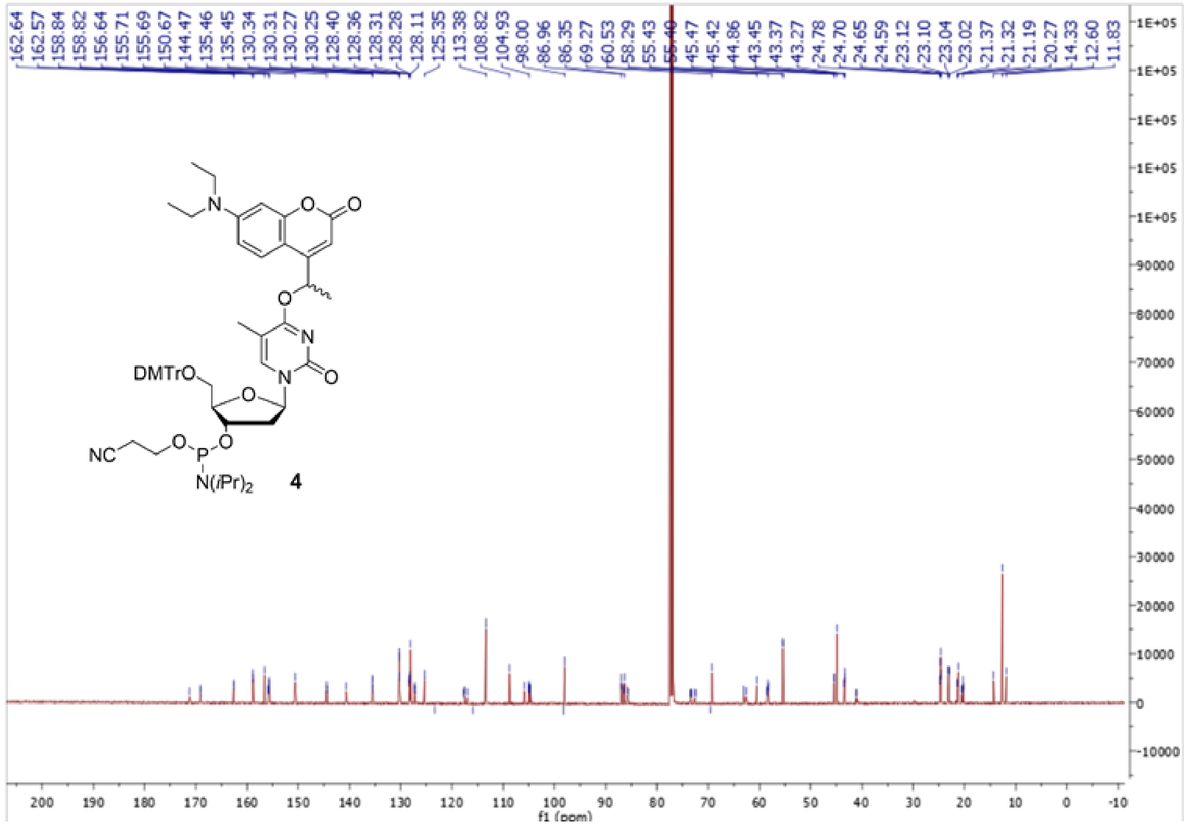

**Figure S20.** $^{13}C$ NMR of **4**.

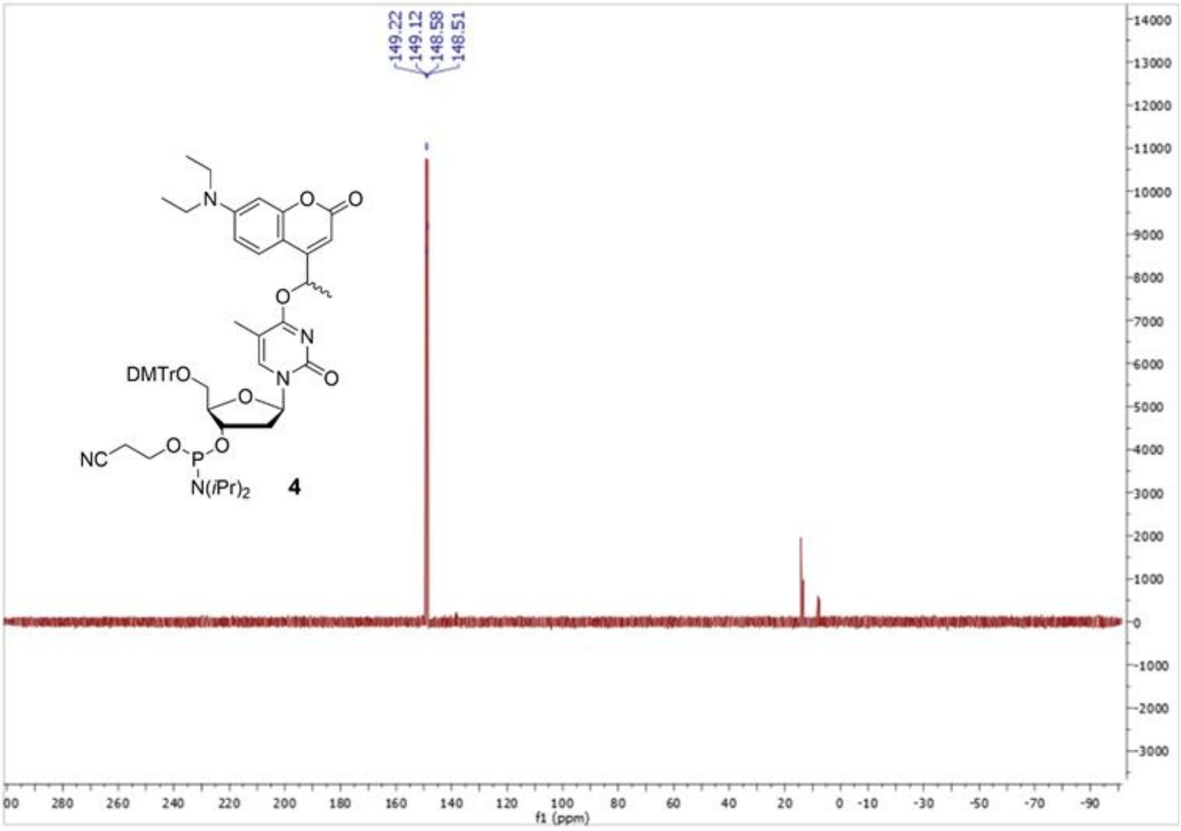


**Figure S21.** $^{31}P$ NMR of **4**.

### Mass Spectra

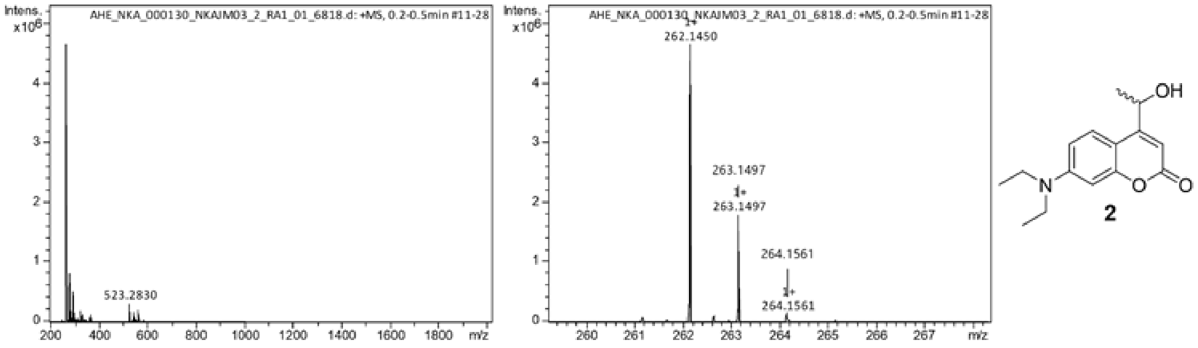


**Figure S22.** High-resolution mass spectrum of **2**.

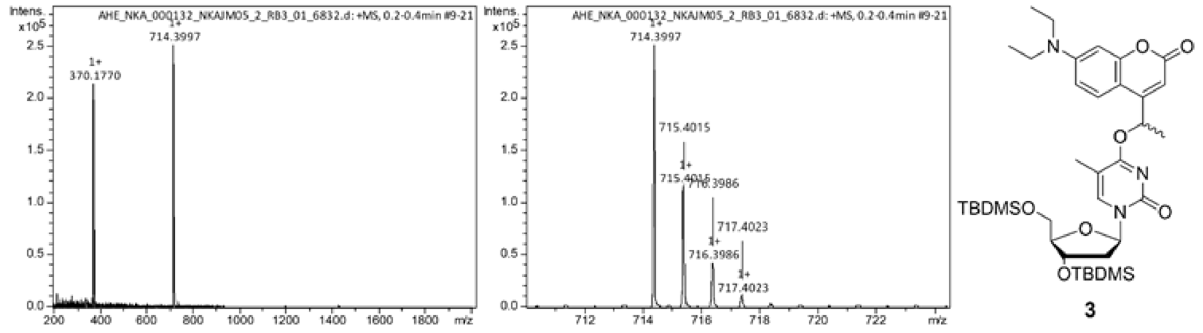


**Figure S23.** High-resolution mass spectrum of **3**.

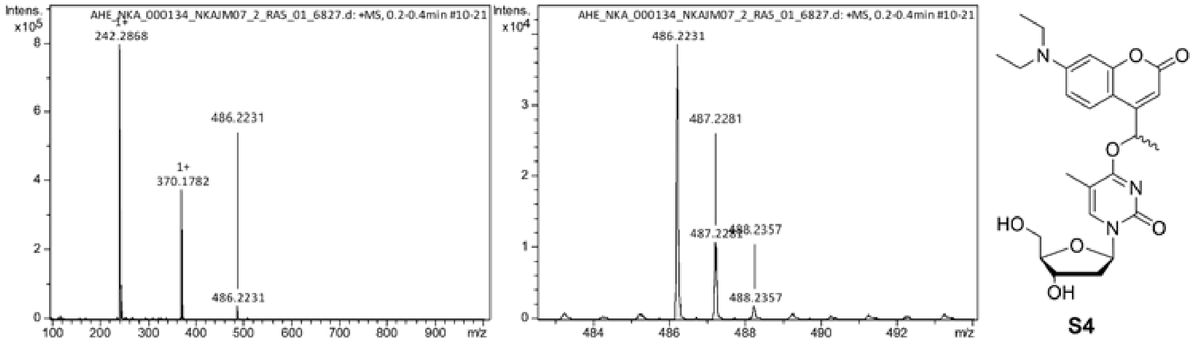


**Figure S24.** High-resolution mass spectrum of **S4**.

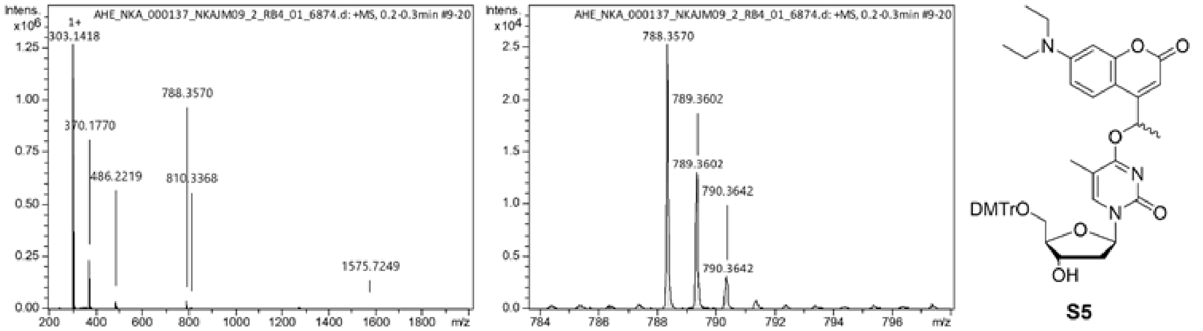


**Figure S25.** High-resolution mass spectrum of **S5**.

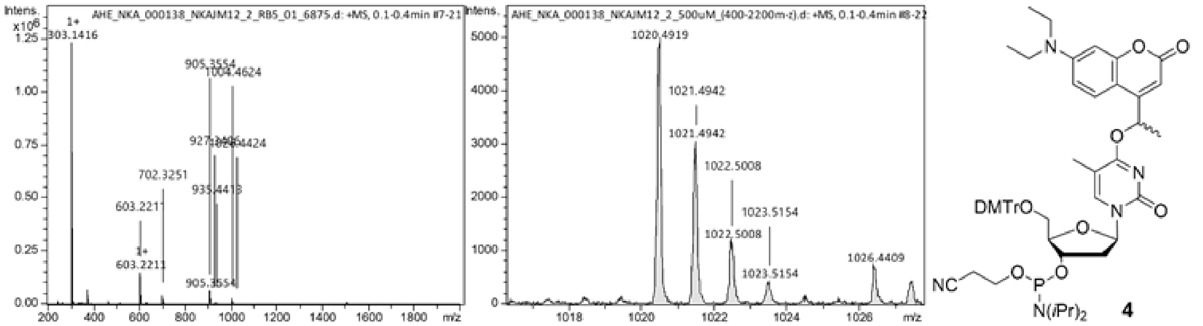


**Figure S26.** High-resolution mass spectrum of **4**.

#### Oligonucleotide Mass Spectra

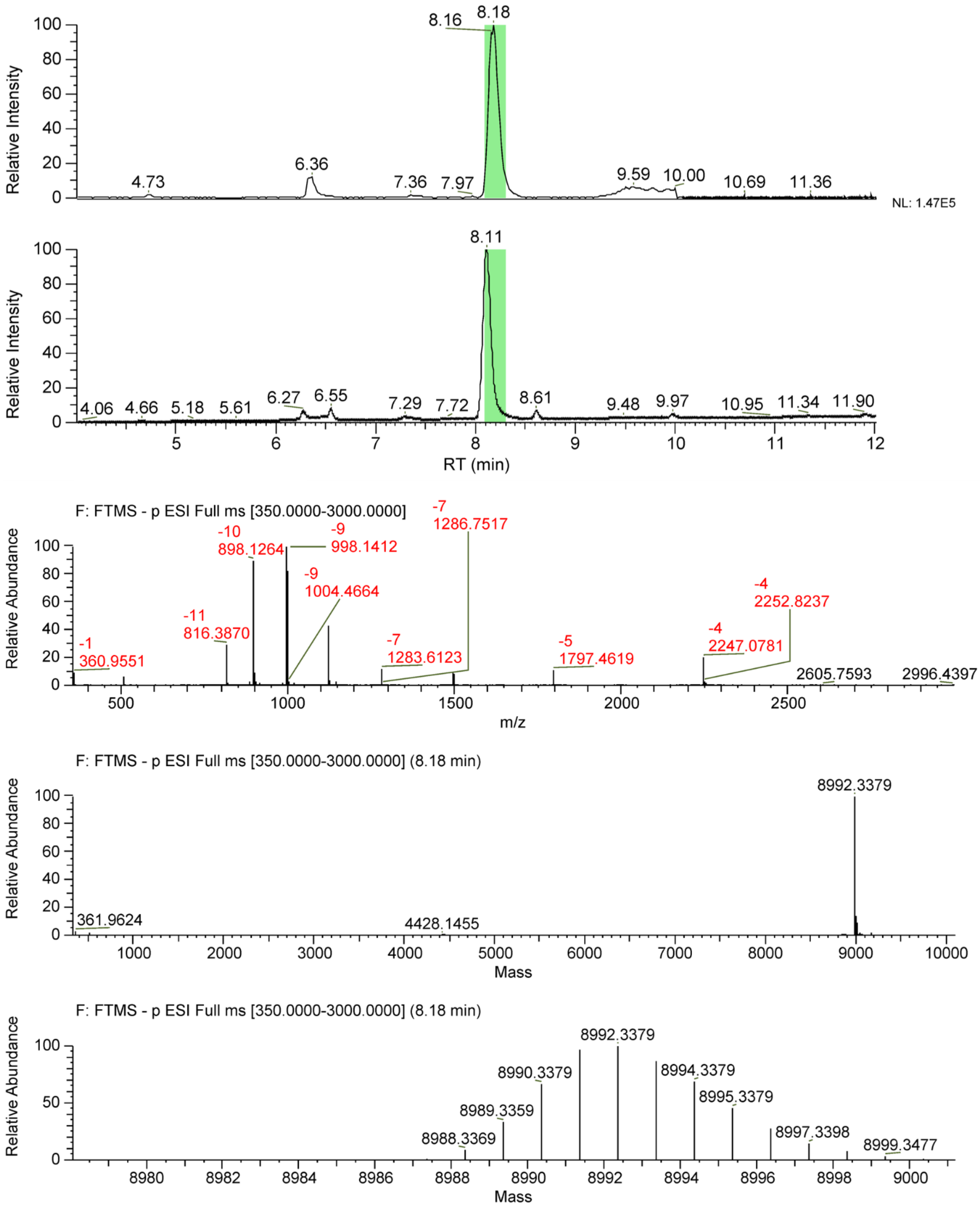


**Figure S27.** Mass spectrometry characterization of **cP1a**.

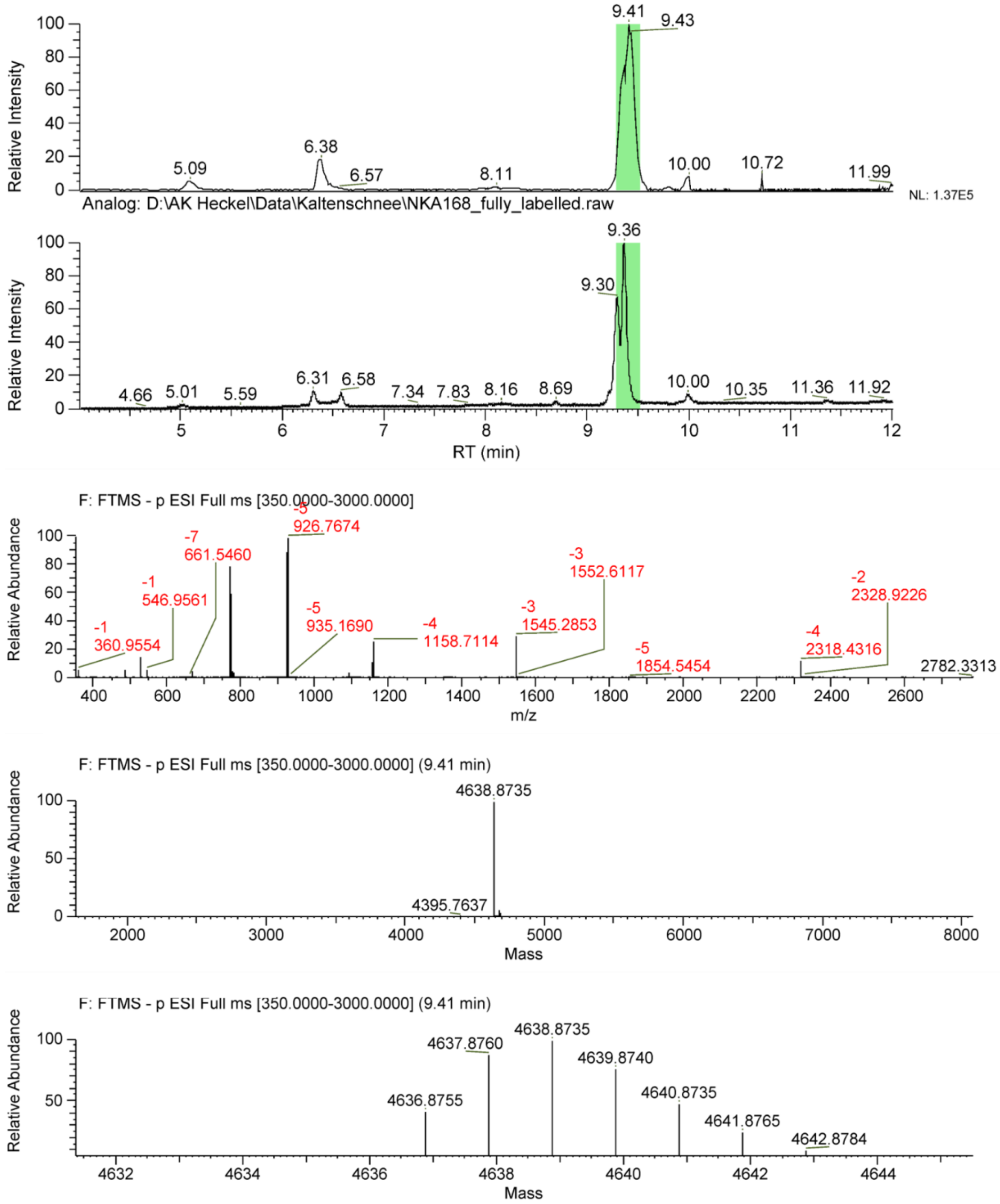


**Figure S28.** Mass spectrometry characterization of **cP1$_b$**.

### References


[1] P. Seyfried, L. Eiden, N. Grebenovsky, G. Mayer, A. Heckel, *Angew Chem Int Ed Engl* **2017**, *56*, 359–363.

[2] J. Schnitzbauer, M. T. Strauss, T. Schlichthaerle, F. Schueder, R. Jungmann, *Nat Protoc* **2017**, *12*, 1198–1228.

[3] J. Vogelsang, R. Kasper, C. Steinhauer, B. Person, M. Heilemann, M. Sauer, P. Tinnefeld, *Angew Chem Int Ed Engl* **2008**, *47*, 5465–5469.

[4] P. Blumhardt, J. Stein, J. Mücksch, F. Stehr, J. Bauer, R. Jungmann, P. Schwille, *Molecules* **2018**, *23*, DOI 10.3390/molecules23123165.

[5] A. D. Edelstein, M. A. Tsuchida, N. Amodaj, H. Pinkard, R. D. Vale, N. Stuurman, *J Biol Methods* **2014**, *1*, DOI 10.14440/jbm.2014.36.

[6] U. Endesfelder, S. Malkusch, F. Fricke, M. Heilemann, *Histochem Cell Biol* **2014**, *141*, 629–638.
